# Activation of Disease-Associated Polycystin-1 Mutants by Binding of Stalk-derived Agonist Peptides

**DOI:** 10.64898/2026.10.10.758211

**Authors:** Keya Joshi, Brenda S. Magenheimer, Allan Haldane, Yan Zhang, Robin L. Maser, Yinglong Miao

**Affiliations:** Department of Pharmacology and Computational Medicine Program, University of North Carolina – Chapel Hill, Chapel Hill, NC, United States, 27599; Bioinformatics and Computational Biology Program, University of North Carolina – Chapel Hill, Chapel Hill, NC, United States, 27599; Departments of Biochemistry and Molecular Biology; Clinical Laboratory Sciences; The Jared Grantham Kidney Institute, University of Kansas Medical Center, Kansas City, KS, United States, 66160; Department of Physics, and Center for Biophysics and Computational Biology, Temple University, Philadelphia, PA, United States, 19122; Department of Biological Sciences, Michigan Technological University, Houghton, Michigan, United States, 49931

## Abstract

Polycystin-1 (PC1) functions as an atypical G-protein-coupled receptor (GPCR) with an autocatalytic GAIN domain that cleaves PC1 into extracellular N-terminal (NTF) and membrane-embedded C-terminal (CTF) fragments. Mutations in PC1 are responsible for 85% of autosomal dominant polycystic kidney disease (ADPKD) cases. Activation of signaling by PC1 CTF has been shown to be allosterically mediated by a stalk tethered agonist (TA) and to be prevented by mutations of the stalk. Soluble, synthetic peptides derived from the PC1 stalk were also shown to re-activate signaling by a stalkless CTF mutant. Here, we have investigated the ability of selected stalk peptide agonists p9, p15 or p17 to re-activate signaling by PC1 CTF with ADPKD-associated missense mutations, G3052R or R3063C. Biochemical experimental data showed that each of the three stalk peptides was able to significantly reactivate signaling by the CTF-R3063C mutant. In contrast, signaling by CTF-G3052R could only be rescued by treatment with p17. In parallel, we employed peptide docking and novel Peptide Gaussian-accelerated Molecular Dynamics (Pep-GaMD) simulations to elucidate the binding conformations of p9, p15 and p17 to the PC1 mutants. The simulation findings were highly consistent with experimental data. Furthermore, Pep-GaMD revealed binding conformations of the agonist peptides and important low-energy conformational states of the PC1 CTF, as well as their critical residue interactions. Overall, our complementary approach provides important mechanistic insights into the re-activation of PC1 disease-associated mutants by stalk-derived peptide agonists. Such insights are expected to be valuable for the future design and development of peptide modulators targeting PC1 for an effective ADPKD therapeutic treatment.

**Visual Abstract:** 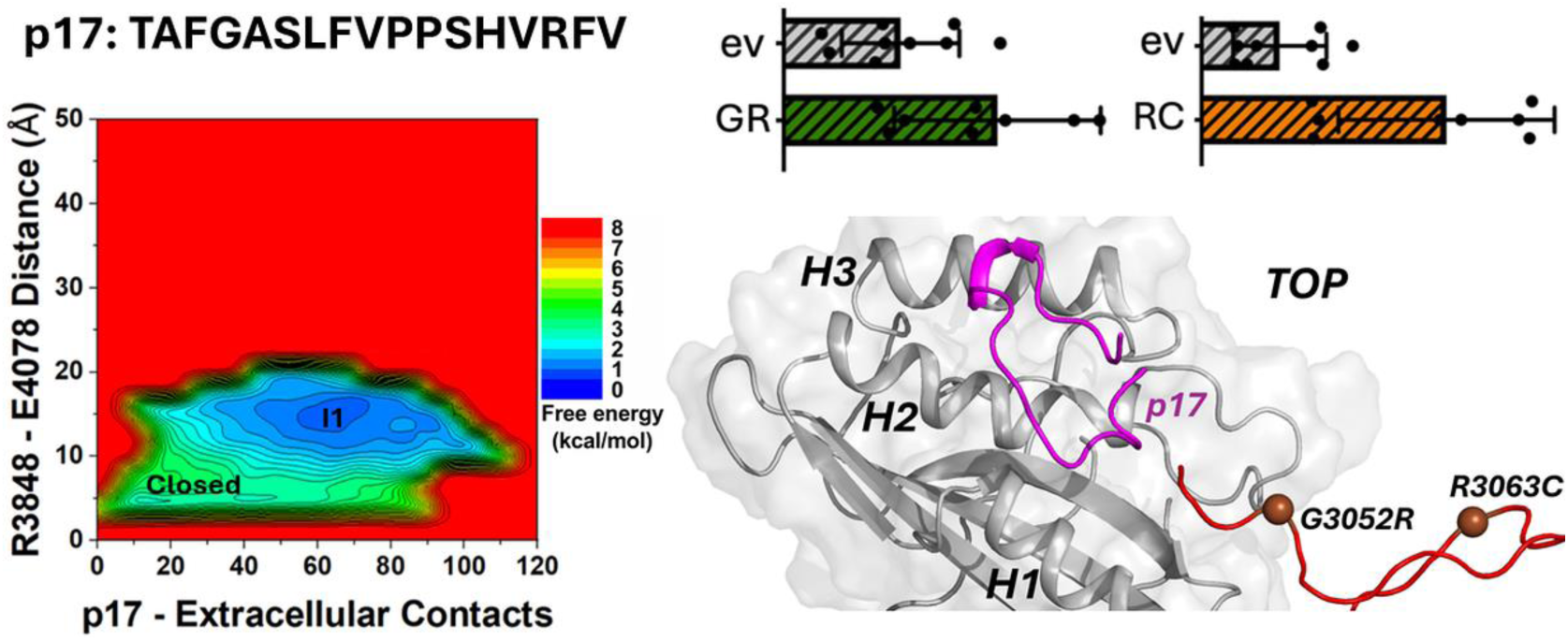

Structural dynamic models are presented for binding of novel synthetic, soluble peptide agonists and associated activation of disease-associated Polycystin-1 mutants through a combination of complementary cellular signaling assays and accelerated molecular simulations. The p17 peptide derived from the first 17 residues of the protein stalk tethered agonist is shown here.

## 1. INTRODUCTION

Autosomal dominant polycystic kidney disease (ADPKD) is the most common monogenetic disorder that manifests in renal disease, affecting nearly 6 million people worldwide^1, 2^. Mutations in two genes, *PKD1* and *PKD2*, whose products are the polycystin-1 (PC1) and polycystin-2 (PC2) proteins, are the major cause of ADPKD^3–5^. The hallmark of ADPKD is the development of bilateral, fluid-filled, renal cysts, which increase in size and number with age and lead to renal failure in half of the cases^3–5^. PC1 functions as an atypical G-protein-coupled receptor (GPCR) and has been implicated in the regulation of cytosolic cyclic adenosine monophosphate (cAMP) concentrations^6, 7^. PC2 functions as a non-specific cation channel and can alter intracellular Ca^2+^ concentrations^8, 9^. In addition, PC1 and PC2 proteins have been known to interact and are currently thought to form a signaling receptor-ion channel complex^10–14^. In 2018, the structure of the human PC1–PC2 complex (RCSB PDB: 6A70) was resolved as a heterotetramer consisting of one PC1 and three PC2 protein subunits^15^ in which the last two transmembrane (TM) domains of each protein form the ion channel pore^16^, which is surrounded by the large, extracellular TOP (Tetragonal Opening of Polycystins) domain from each subunit.

Currently, the only approved ADPKD treatment is based on a small molecule - Jynarque^TM^ which functions as an antagonist of the vasopressin V2 receptor thus lowering renal cAMP levels^17–19^. However, this treatment has serious side effects and can only reduce the disease progression^20^. A significant portion of *PKD1* mutations is caused by non-truncating mutations that affect the production of mature PC1 protein or produce mutant PC1 protein with reduced function^21–23^. Numerous strategies are being pursued to increase the amount of residual, functional PC1 protein^24–26^. As an alternative but related approach, we have been exploring the development of small molecules or peptides that directly target and activate (or augment) the signaling function of PC1.

PC1 is composed of 11 TM domains and has a short C-terminal cytosolic tail (C-tail)^27, 28^. The C-tail is involved in binding and activating G-protein signaling^29–33^ while the protein’s N-terminal, extracellular region consists of multiple binding domains that mediate cell-to-cell and cell-to-matrix interactions and bind various Wnt ligands^14, 34–37^. PC1 and the Adhesion family of GPCRs (ADGRs) share a conserved GPCR proteolysis site (GPS) near their first TM domain that undergoes self-catalyzed cleavage resulting in an N-terminal, extracellular fragment (NTF) and a C-terminal, membrane-embedded fragment (CTF) that remain noncovalently attached after cleavage^38–42^. Following GPS cleavage and removal of the NTF, a short, extracellular “stalk” at the N-terminus of the CTF becomes exposed which further binds and interacts with the seven TM domains causing conformational changes that activate G-protein signaling, i.e., the tethered agonist (TA) mechanism^43–49^. In addition, studies with numerous ADGRs have shown that synthetic peptides derived from the stalk sequence can also function as soluble agonists to initiate signaling activation^41, 45^.

Recently, we have shown that both the tethered stalk and synthetic, stalk-derived peptides of PC1 interact with its TOP domain and mediate signaling activation^50, 51^. The lid-like TOP domain is composed of three α-helices (*H1*-*H3*), five anti-parallel β-sheets (*β1*-*β5*), and two extended interdomain interaction motifs called Finger-1 and Finger-2 **(Figure S1A)**^15, 52^. The model we proposed for the tethered stalk-mediated activation of PC1 represents an allosteric mechanism that involves the formation of a key salt bridge between residue R3848 from the TOP domain and residue E4078 in the putative pore loop (PL) along with a stable hydrogen-bonded residue-residue interaction between N3074 at the base of the stalk and S3585 near the extracellular end of the fifth

TM domain. Our previous analysis of the ability of stalk-derived peptides to activate signaling of PC1 involved a mutant CTF lacking the stalk domain (CTF-Δstalk)^50^ and revealed specific binding regions of the peptide agonists using the recently developed Peptide Gaussian-accelerated Molecular Dynamics (Pep-GaMD) method^53^. In Pep-GaMD, the essential potential energy of the peptide is selectively boosted to account for the peptide’s high flexibility, which enhances the peptide-protein interactions. In addition, another boost-potential is applied to the remaining potential energy of the entire system in a dual boost algorithm to enhance the peptide diffusion and protein conformational changes. Moreover, because the boost potential exhibits a near-Gaussian distribution, free energy profiles can be properly recovered through cumulant expansion to the second order^54, 55^. Both previous studies^50, 51^ identified the binding interface for the tethered stalk and the stalk-derived peptide agonists to be within the *H1* and *H2* helices and the β3, β4, and β5 strands of the PC1 TOP domain^50^.

In this study, we have combined cellular signaling assays to test the ability of selected stalk-derived peptide agonists to rescue activation by ADPKD-associated mutants of PC1 with *in silico* studies to uncover mechanisms of peptide binding and signaling rescue of the mutant proteins. Synthetic peptides of 9, 15 and 17 residues in length derived from the N-terminus of the PC1 CTF stalk sequence were tested with two signaling-dead missense mutants, G3052R and R3063C, of PC1 CTF. We found that each of the three stalk peptides was able to significantly activate signaling by the CTF-R3063C mutant while in contrast, signaling by CTF-G3052R could only be rescued by treatment with p17. All-atom Pep-GaMD simulations^53^ with the peptide agonists and CTF mutants refined the docking conformation of the peptides bound to a site consisting primarily of the first two alpha helices (*H1* and *H2*) within the extracellular TOP domain of PC1. Key residue-residue interactions within the CTF mutants, including the salt bridge between R3848-E4078 and polar interaction between N3074-S3585, were observed for the binding of peptides capable of re-activating the PC1 CTF mutants, thus demonstrating the consistency between *in vitro* and *in silico* results.

This work has shown the ability of stalk-derived peptide agonists to rescue the signaling activation of two ADPKD-associated missense mutants of PC1. Importantly, Pep-GaMD simulations have provided mechanistic insights into the activation-inducing binding mechanisms of the peptide agonists to PC1 CTF-G3052R and CTF-R3063C disease-associated mutants.

## 2. METHODOLOGY

### 2.1 DNA expression constructs, antibodies and peptide synthesis

Expression constructs of human PC1 mutant CTF-Δstalk, CTF-G3052R and CTF-R3063C were made in pCI vector (Promega) as described previously^51^. Primary antibodies used included E8, for detection of PC1 CTF (PKD RRC) and beta-actin (Sigma). Secondary antibody-HRP conjugates were purchased from Sigma or Jackson ImmunoResearch. Stalk-derived peptides were synthesized using the Fmoc method and verified by HPLC-MS analysis (GenScript).

### 2.2 Cell culture, transient transfection, stalk treatment and luciferase assay

HEK293T cells (ATCC) were maintained and transiently transfected by calcium phosphate method as described previously^51^. Cells (150,000 cells/well) were plated into 24-well plates (3 wells per transfection condition) and transfected with a DNA mixture containing the pNFATluc (firefly luciferase) reporter (100 ng; Stratagene), pGL4.70 [hRluc] (*Renilla* luciferase; 50 ng; Promega), and expression vector encoding mutant CTF constructs (75 ng/3 wells) or an equimolar amount of empty expression vector (ev). Total DNA amount was filled with pBlueScript (Stratagene) to 8 ug. Two hours following medium exchange, one-half of the culture medium volume was replaced with an equal volume of either serum-free medium (no peptide control) or stalk peptide p9, p15 or p17 (2 mM; final concentration of 1 mM) and incubation was continued overnight. Cells were lysed in 1 X Passive Lysis buffer (Promega) with protease and phosphatase inhibitors at 24 hr following the initial peptide or control medium addition. Luciferase activities were determined using the Dual Luciferase Assay Kit (Promega). Firefly luminescence was normalized to Renilla luminescence for each well within a treatment condition (n=3 wells/condition), averaged together for the 3 experiments, and were graphed after setting each value relative to its empty expression vector (ev) control condition. Experiments were performed a minimum of three times (i.e., >3 biological replicates) each with three technical replicates/conditions except where noted.

### 2.3 Western blot analyses

Gel-loaded volumes of cell lysates were calculated based on normalization to relative Renilla luciferase activity values within a condition. Denaturing 7.5% polyacrylamide gels were transferred to nitrocellulose with the TransBlot Turbo and ReadyBlot transfer buffer (BioRad). Blots were incubated with E8 antibody (1:2000) in TBST (10 mM Tris pH 7.4, 150 mM NaCl, 0.1% Tween-0) and 5% powdered dry milk overnight at 4° C. Secondary antibody-HRP (1:5,000) was incubated in TBST/5% milk for 1 hr at room temperature. Blots were developed with Clarity chemiluminescent substrate (BioRad). Multiple exposures were captured for each blot with the Amersham Imager 600 with the band saturation detection mode enabled. Blots were stripped and reprobed with anti-actin (1:1,000) as above.

### 2.4 Statistical analysis

Analyses were performed with GraphPad Prism 9 (GraphPad Software, San Diego, CA, USA). Data are presented as means ± standard deviation. Multiple comparisons used one-way ANOVA and Tukey’s post-hoc analysis. p≤0.05 was considered statistically significant.

### 2.5 Peptide Docking

Computational model of wild type PC1 CTF was obtained from our previous study^51^ in which the PC1 protein subunit had been extracted from the cryo-EM structure of the PC1–PC2 complex (RCSB PDB: 6A70)^15^, and missing regions, including the stalk, PL, and C-tail had been added^51^. The HPEPDOCK webserver^56^ was applied to dock the p9, p15, and p17 stalk-derived peptide agonists to our wild type PC1 CTF computational model **(Figure S2A,B and S3A,B)**.

A total of 10 docking poses were obtained for each peptide agonist, and the top-ranked pose was selected for each peptide based on the docking score and a docking pose fulfilling the following criteria: (i) localization of the peptide within the extracellular TOP domain of PC1 CTF, which excluded poses bound to the intracellular region, PLAT domain, PL, or TM region; and (ii) binding to the previously identified binding region comprised of the *H1* and *H2* helices and adjacent β3– β5 sheets of the TOP domain^50, 51^ **(Figure S1B)**.

All three peptide agonists (p9, p15 and p17) bound primarily to the *H1*-*H2* region in the TOP domain **(Figure S2B and S3B)**. CHARMM-GUI^57^ mutation function was used to apply the ADPKD missense mutation (G3052R or R3063C) within the CTF stalk and refinement of the peptide docking poses was performed using Pep-GaMD simulations^53^.

### 2.6 Pep-GaMD Simulations

The CTF-G3052R and CTF-R3063C computational models were embedded in a palmitoyl-oleoyl-phosphatidyl-choline (POPC) bilayer and solvated in 0.15 M NaCl explicit solvent using CHARMM-GUI webserver^57^ **(Figure S2C and S3C)**. Neutral patches (acetyl and methylamide) were added to the terminal residues of the protein, while peptide termini were kept as charged (*NH*^+^ and COO^−^). The CTF-G3052R simulation systems measured ∼135×135×171 Å^3^ in dimension, containing a total of ∼300,000 atoms with explicit solvent and lipid molecules **(Table S1 to S3)**. The CTF-R3063C simulation systems measured ∼136×136×171 Å^3^ in dimension, containing a total of ∼300,000 atoms with explicit solvent and lipid molecules **(Table S4 to S6)**. The CHARMM36m force field parameters^58, 59^ were used for the protein, peptides, and lipids. CHARMM-GUI^57^ output files and scripts were used with default parameters to prepare the systems for Pep-GaMD simulations^53^.

Energy minimization was performed for 5000 steps using a constant number, volume, and temperature (NVT) ensemble at 310 K. Further equilibration was done for 375 ps at 310 K using a constant number, pressure, and temperature (NPT) ensemble. Conventional MD simulations were performed on the systems for 40 ns at 1 atm pressure and 310 K temperature. All-atom Pep-GaMD^53^ simulations were performed with a short cMD for 12 ns, Pep-GaMD^53^ equilibration for 48 ns and five independent Pep-GaMD^53^ production runs for 1000 ns on each system with randomized initial atomic velocities. A cutoff distance of 9 Å was used for the van der Waals and short-range electrostatic interactions. Long-range electrostatic interactions were computed with the particle-mesh Ewald summation method^60^.

### 2.7 Simulation Analysis

Pep-GaMD simulation trajectories were analyzed using CPPTRAJ^61^ and VMD^62^ tools. Trajectory analysis showed peptides interacting with residues in the TOP domain of CTF-G3052R and CTF-R3063C. A previously identified salt bridge formed between the TOP domain residue R3848 and the pore loop residue E4078 of the PC1 CTF protein is a characteristic feature of its active, closed conformation state^50, 51^. The number of residue contacts formed between each peptide and the extracellular domains of PC1 CTF **(Figure S4)**, and the R3848-E4078 salt bridge distance **(Figure S5)** were used as reaction coordinates to calculate the 2D free energy profiles using the PyReweighting toolkit^55^. Residue contacts were calculated for heavy atoms with any atom pair having a distance less than or equal to 4 Å. A bin size of 1 Å was used for the distances and 10 for the number of contacts.

The angle between the principal axes of *H1*-*H2* helices of the TOP domain was used to characterize the peptide agonists’ binding site (determined in previous study)^50^. For each simulation frame, the principal axis vector of each helix was obtained by principal component analysis of the corresponding Cα coordinates. The *H1*–*H2* inter-helical angle (θ) was then calculated from the dot product of the two normalized axis vectors *θ* = *cos*^−1^(|v₁ · v₂|), where v₁ and v₂ represent the principal axis vectors of *H1* and *H2* helices, respectively.

The five independent 1000 ns Pep-GaMD^53^ production simulations for each system were combined to perform structural clustering using the dpeak clustering algorithm^63^ in CPPTRAJ. PyReweighting toolkit^55^ was then applied to calculate the original free energy values of each peptide structural cluster with a cutoff of 500 frames. The structural clusters were finally ranked according to the reweighted free energy values. In the subsequent results, peptide residues are numbered relative to the N-terminus of the stalk starting with 1, while residues of CTF-G3052R and CTF-R3063C are numbered relative to the GPS cleavage site of human PC1 CTF that starts with T3049.

## 3. RESULTS AND DISCUSSION

### 3.1 Stalk-derived peptide agonists p9, p15 and p17 differentially activated disease-associated mutants of PC1 CTF

Our previous work showed that deletion of the N-terminal-most 21-amino acid residues of the PC1 CTF stalk (CTF-Δstalk; **Figure 1A**) prevents CTF-mediated activation of a pNFATluc reporter^51^. In addition, soluble synthetic peptides derived from the PC1 stalk sequence were able to stimulate signaling by the CTF-Δstalk mutant^50^, thus providing further support for the stalk region of PC1 CTF functioning as a tethered peptide agonist. Here, we investigated whether selected stalk-derived peptide agonists could rescue signaling by CTF proteins with an ADPKD-associated missense mutation, namely G3052R and R3063C. We chose CTF-G3052R and CTF-R3063C for our investigation because (i) the variant residues are located within the critical 21-amino acid region of the PC1 stalk tethered agonist (TA) **(Figure 1A)**, (ii) these constructs were previously found to be defective in cellular signaling assays, and (iii) their computer models were rarely observed to exhibit the *in silico* residue-residue interactions characteristic of activated wild-type CTF^51^.

**Figure 1.**
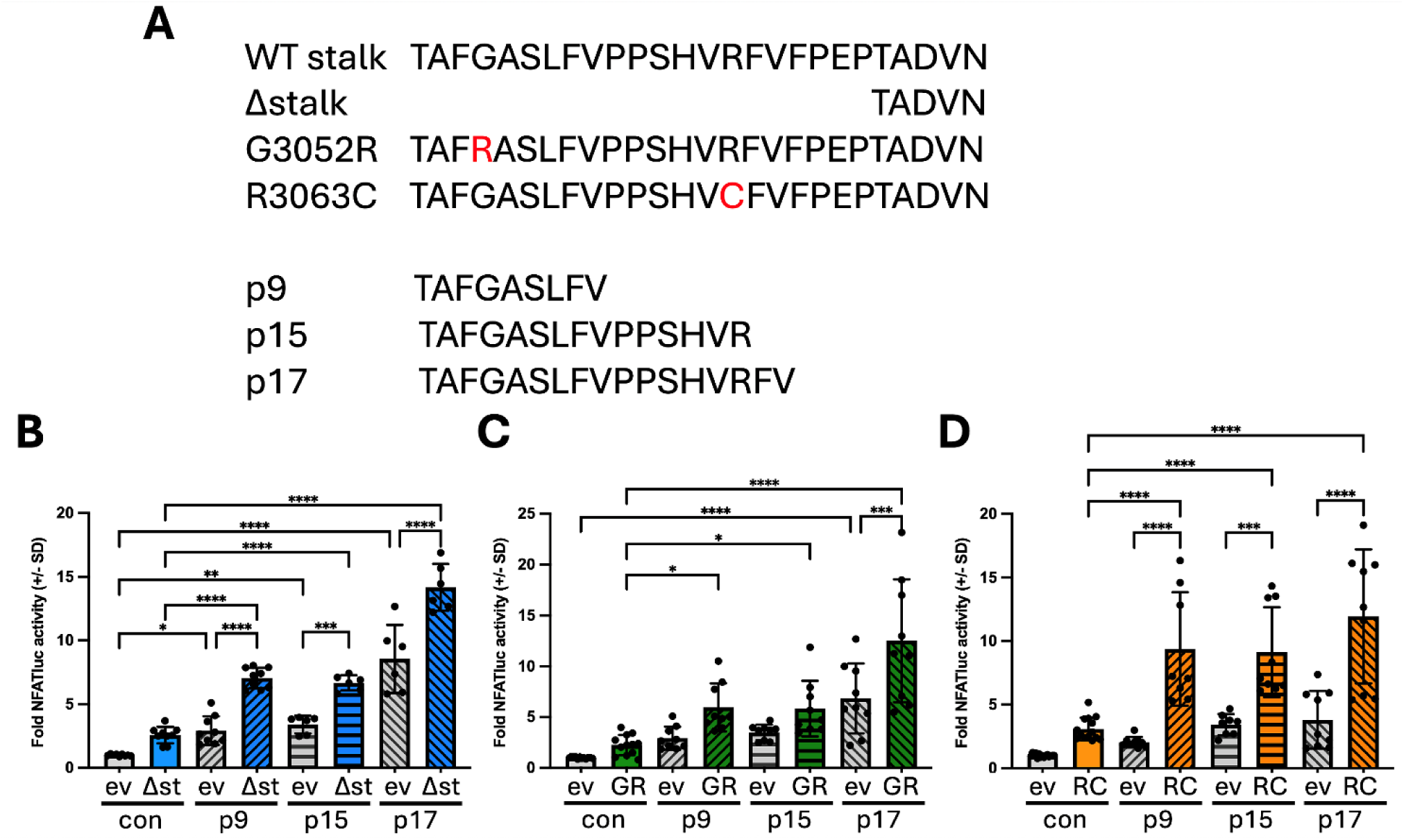
Activation of PC1 CTF stalk mutants by synthetic stalk-derived peptides: **(A)** Alignment of stalk sequences for wild type (WT) and mutant human PC1 CTF expression constructs (top) and synthetic stalk-derived peptides (bottom). **(B-D)** Stalk peptide treatment of empty expression vector (ev)-or mutant stalk CTF-transfected cells. Results of pNFAT-luc reporter activation by ev-(gray bars) and CTF Δstalk (Δst, blue bars) **(B)**, CTF-G3052R (GR, green bars) **(C)**, or CTF-R3063C (RC, orange bars) **(D**) expression constructs after 24 hr treatment with (hatched bars) or without (con, solid bars) stalk peptide p9, p15 or p17 (hatched bars). Each graph shows fold reporter activation relative to the ev control condition. Each bar represents the mean (+/-SD) of three separate experiments, each consisting of 2-3 wells/condition. *p<0.05; **p<0.01; ***p=0.001; ****p<0.0001. Analysis by one-way ANOVA with Sidak’s Multiple Comparisons post-test.

Stalk-derived peptide agonists consisting of the first 9-, 15-, or 17-residues (i.e., p9, p15, p17) **(Figure 1A)** from the PC1 stalk TA sequence were selected for analysis based on their differential re-activation abilities with the stalkless CTF mutant (CTF-Δstalk)^51^. HEK293T cells were transfected with the pNFATluc reporter and either empty expression vector (ev), or expression constructs encoding CTF-Δstalk, CTF-G3052R or CTF-R3063C and were treated either with culture medium alone (i.e., no peptide control) or with one of the stalk peptide agonists p9, p15, or p17 **(Figure 1A)**.

As anticipated, none of the CTF stalk mutants could activate the NFAT reporter under control conditions (i.e., without peptide treatment) **(Figure 1B-D)**. However, treatment with each of the 3 stalk peptides was able to significantly reactivate signaling by both CTF-Δstalk and CTF-R3063C mutants to the reporter when compared to both ‘background’ control conditions, i.e., ev + peptide and mutant control **(Figure 1B, D)**. In contrast, neither p9 nor p15 were able to rescue CTF-G3052R signaling to the reporter; only treatment with p17 was capable of reactivation of this mutant **(Figure 1C)**. Western blot analysis showed expression levels of CTF-G3052R and - R3063C mutants were comparable to CTF-Δstalk in both control- and peptide-treated conditions **(Figure S6)**, thus eliminating differences in protein expression or stability as responsible for the inability of p9 or p15 to re-activate signaling by CTF-G3052R. Previous work had shown that all three CTF mutants were able to traffic to the cell surface^51^. These experimental results thus suggested that the ability of different stalk-derived peptide agonists to rescue the allosteric activation of PC1 CTF is influenced by its specific stalk mutation.

### 3.2 Binding of p9 or p15 does not induce the active conformation of CTF-G3052R

To provide mechanistic insight into the differential rescue of signaling activation by the selected peptide agonists, we combined peptide docking and Pep-GaMD simulations to examine binding of the p9, p15, and p17 peptides to the CTF-G3052R and CTF-R3063C mutants. Through peptide docking using HPEPDOCK, all three peptides were found to bind the protein surface of the TOP domain between the *H1* and *H2* helices **(Figure S2B and S3B)**. Refinement of the peptide docking poses was performed using Pep-GaMD simulations.

For the p9-bound CTF-G3052R, a 2D free energy profile calculated from reweighting of five 1000ns Pep-GaMD simulations allowed us to identify three low-energy states designated as ‘Open’, ‘Intermediate 1’ **(I1)** and ‘Intermediate 2’ **(I2) (Figure 2A)**. In the **Open** state, although p9 remained bound to the *H1*-*H2* region, the important “signaling activation interaction” between TOP domain residue R3848 and PL residue E4078^50, 51^ remained quite distant from each other at ∼34 Å **(Figure 2A)**. In the **I1** state, characterized by a ∼16 Å distance between residues R3848 and E4078 **(Figure 3B)**, p9 retained a linear conformation and remained bound to the *H1*-*H2* helices in the TOP domain **(Figure 3A)** and formed two hydrogen-bonded interactions between peptide-protein residue pairs V9-K3706 and S6-H3729, and one hydrophobic interaction between the F8-P3733 residue pair **(Figure 3A)**. In the **I2** state, characterized by a ∼26 Å distance between residues R3848 and E4078 **(Figure 3D)**, p9 remained bound between the helices *H1* and *H2* and retained a linear conformation similar to the **I1** state **(Figure 3C)**. Two hydrogen-bonded interactions were also observed between the following peptide-protein residue pairs S6-Q3702 and V9-K3706 **(Figure 3C)**. The hydrogen-bonded interaction between stalk residue N3074 and TM5 residue S3585, previously identified to be a feature of the signaling-active WT PC1 CTF^51^, maintained a distance around ∼9 Å **(Figure S7A)**, corresponding to an open/inactive state of PC1 CTF protein **(Figure S8A)**.

**Figure 2.**
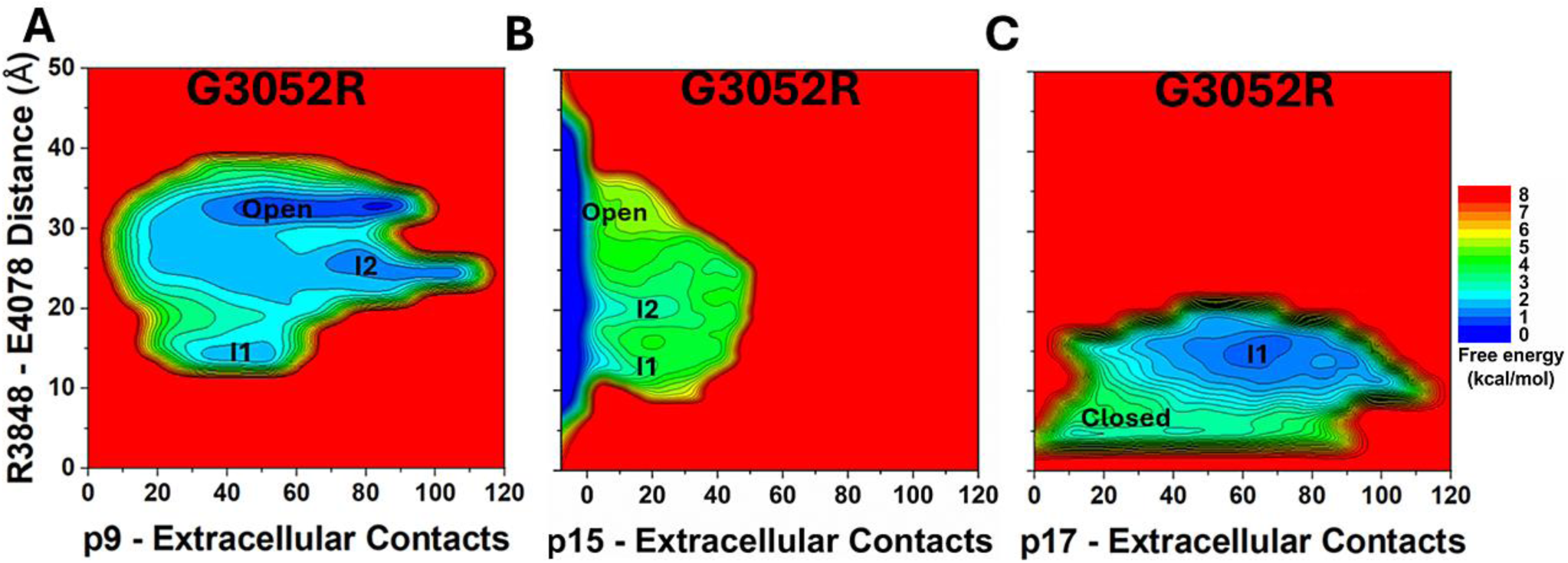
Free energy profiles of p9, p15 and p17 binding to CTF-G3052R calculated from five 1000ns Pep-GaMD simulations on each system: **(A-C)** Free energy profiles of the **(A)** p9-bound, **(B)** p15-bound and **(C)** p17-bound CTF-G3052R regarding the number of atom contacts between each peptide and extracellular domains of CTF and the distance between the CZ atom of R3848 and CD atom of E4078 in CTF.

**Figure 3.**
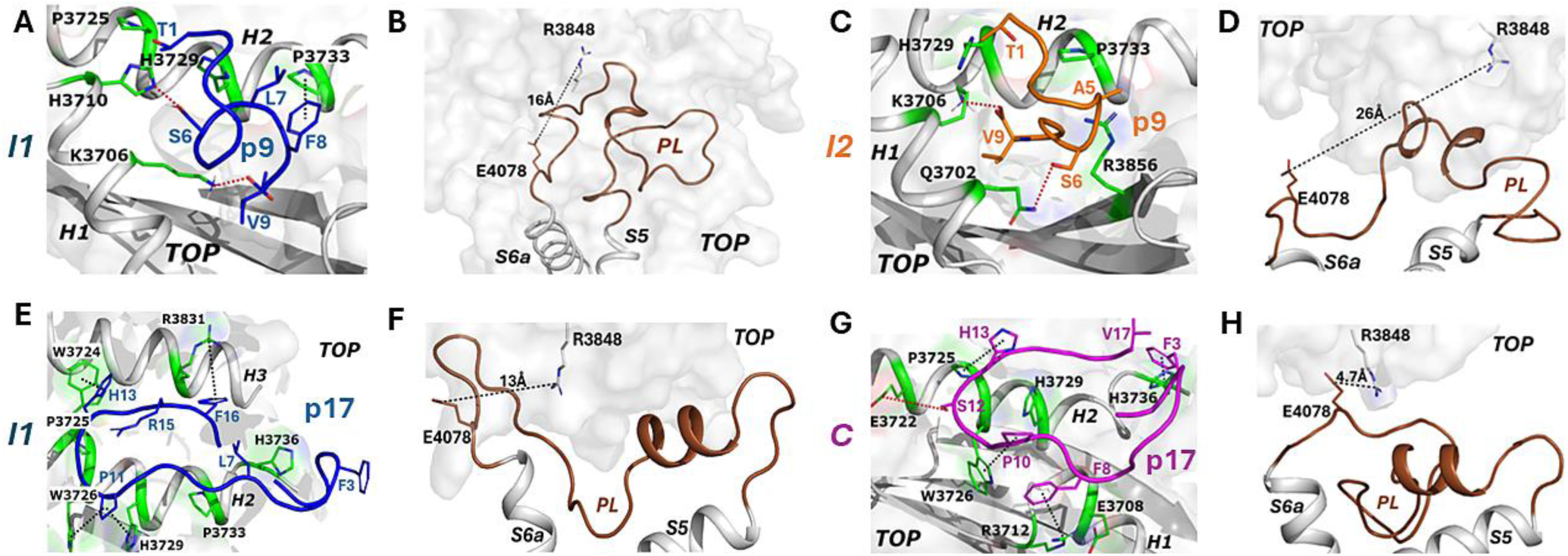
Representative low-energy conformational states of p9 and p17 binding to CTF-G3052R: **(A, C)** Interactions observed between peptide-protein residues in Intermediate I1 and I2 states of p9-bound CTF-G3052R. **(B, D)** Distance between the TOP domain residue R3848 and pore loop (PL) residue E4078 observed in Intermediate I1 and I2 states of p9-bound CTF-G3052R. **(E, G)** Interactions observed between peptide-protein residues in Intermediate I1 and Closed (C) states of p17-bound CTF-G3052R. **(F, H)** Distance between the TOP domain residue R3848 and pore loop (PL) residue E4078 observed in Intermediate I1 and Closed (C) states of p17-bound CTF-G3052R.

Pep-GaMD simulations of the p15-bound CTF-G3052R mutant yielded a free energy profile with three low-energy states designated as ‘Open’, ‘I1’ and ‘I2’ **(Figure 2B)**. In the **Open** state, p15 remained bound to the *H1*-*H2* region, however, the distance between the TOP domain residue R3848 and PL residue E4078 remained quite distant from each other at ∼30 Å **(Figure 2B)**. In the **I1** state, the R3848-E4078 salt bridge interaction was reduced to ∼9 Å **(Figure S9B)**, and p15 retained a linear conformation and became bound between the TOP domain and extracellular loop 1 (ECL1) **(Figure S9A)**. In the **I2** state, characterized by a R3848-E4078 distance of ∼20 Å **(Figure S9D)**, p15 maintained a similar linear conformation as in the I1 state, however, its binding site was shifted below the *H1* helix. One hydrogen-bonded interaction was observed between the peptide-protein residue pair F8-Q3702 **(Figure S9C)**. In both I1 and I2 states, p15 formed extensive hydrophobic contacts with neighboring residues from ECL1 loop and *H1* helix **(Table S7)**. In addition, the distance between the stalk residue N3074 and TM5 residue S3585 remained around ∼9 Å throughout the simulations for the p15-bound CTF-G3035R mutant **(Figure S7B)**, consistent with an open/inactive low energy state **(Figure S8B)**.

### 3.3 Binding of p17 enabled formation of the closed, activated conformation of the CTF-G3052R mutant

Pep-GaMD simulations on the p17-bound CTF-G3052R mutant revealed two low-energy states-‘I1’ and ‘Closed’ **(Figure 2C)**. In the **I1** state, characterized by a R3848-E4078 distance of ∼13 Å **(Figure 3F)**, p17 adopted a circular loop conformation and relocated to the *H3*-*H2* region on the TOP domain **(Figure 3E)** and formed one cation-π interaction between F16-R3831, one hydrophobic interaction between P11-W3726 and two aromatic interactions between H13–W3724 and P11–H3729 peptide-protein residue pairs **(Figure 3E)**.

In the **Closed** state of the p17-bound G3052R mutant, the important signaling activation interaction^50, 51^ between TOP domain residue R3848 and PL residue E4078 was formed at ∼4.7 Å **(Figure 3H)**. p17 remained bound to the TOP domain primarily between the *H1*-*H2* region and adopted a cyclized conformation with the two terminal ends (F3 and V17 residues) coming in close proximity **(Figure 3G)** and formed one hydrogen-bonded interaction between S12-E3722, one cation-π interaction between F8-R3712, one hydrophobic interaction between P10-W3726 and two aromatic interactions between H13-P3725 and F3-H3736 peptide-protein pairs **(Figure 3G)**. In the ‘Closed’ state, the hydrogen-bonded residue-residue interaction between the stalk residue N3074 and TM5 residue S3585 was also observed, consistent with the closed/active low energy state of wild type PC1 CTF **(Figure S7C and S8C)**.

### 3.4 Binding of p9, p15 and p17 enabled formation of the activated conformation of the CTF-R3063C mutant

Free energy profile obtained from Pep-GaMD simulations on the p9-bound CTF-R3063C mutant revealed two low-energy states designated as ‘I2’ and ‘Closed’ **(Figure 4A)**. In the **I2** state, the distance between the TOP domain residue R3848 and PL residue E4078 was centered around ∼16 Å **(Figure S10B)** while p9 retained a linear conformation and remained bound between *H1* and *H2* **(Figure S10A)** on the TOP domain and formed one hydrophobic interaction between F8-W3726, one aromatic interaction between F3-H3710 and one cation-π interaction between F8-R3700 peptide-protein residue pairs **(Figure S10A)**. The **Closed** state of the p9-bound CTF-R3063C mutant was characterized by formation of the salt bridge interaction between TOP domain residue R3848 and PL residue E4078 at ∼4.9 Å **(Figure 5B)**. p9 adopted a cyclized conformation with its two terminal residues, T1 and V9, coming into close proximity and remained primarily bound to the *H1*-*H2* region on the TOP domain **(Figure 5A)** and formed one cation-π interaction between the F3-R3856 peptide-protein pair **(Figure 5A)**. Closed state of p9-bound CTF-R3063C was further characterized by the formation of the hydrogen-bonded interaction between stalk residue N3074 and TM5 residue S3585 consistent with the closed/active state of wild type PC1 CTF protein **(Figure S7D and S8D)**.

**Figure 4.**
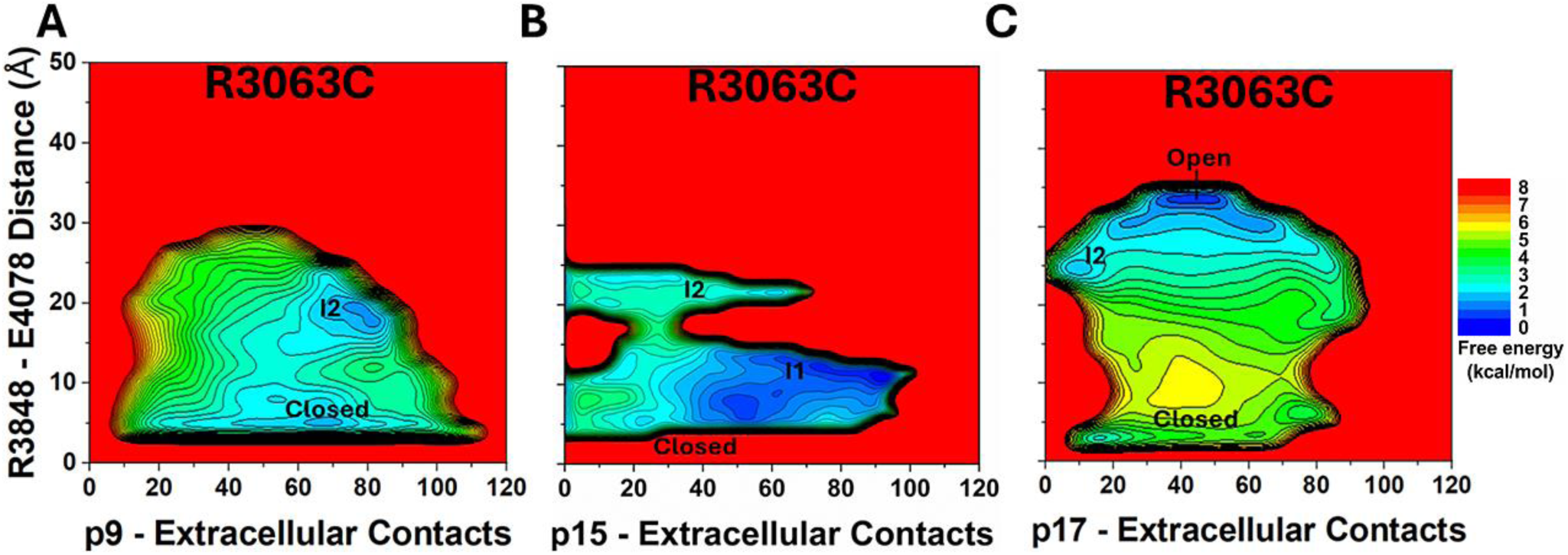
Free energy profiles of p9, p15 and p17 binding to CTF-R3063C calculated from five 1000ns Pep-GaMD simulations on each system: **(A-C)** Free energy profiles of the **(A)** p9-bound, **(B)** p15-bound and **(C)** p17-bound CTF-R3063C regarding the number of atom contacts between each peptide and extracellular domains of CTF and the distance between the CZ atom of R3848 and CD atom of E4078 in CTF.

**Figure 5.**
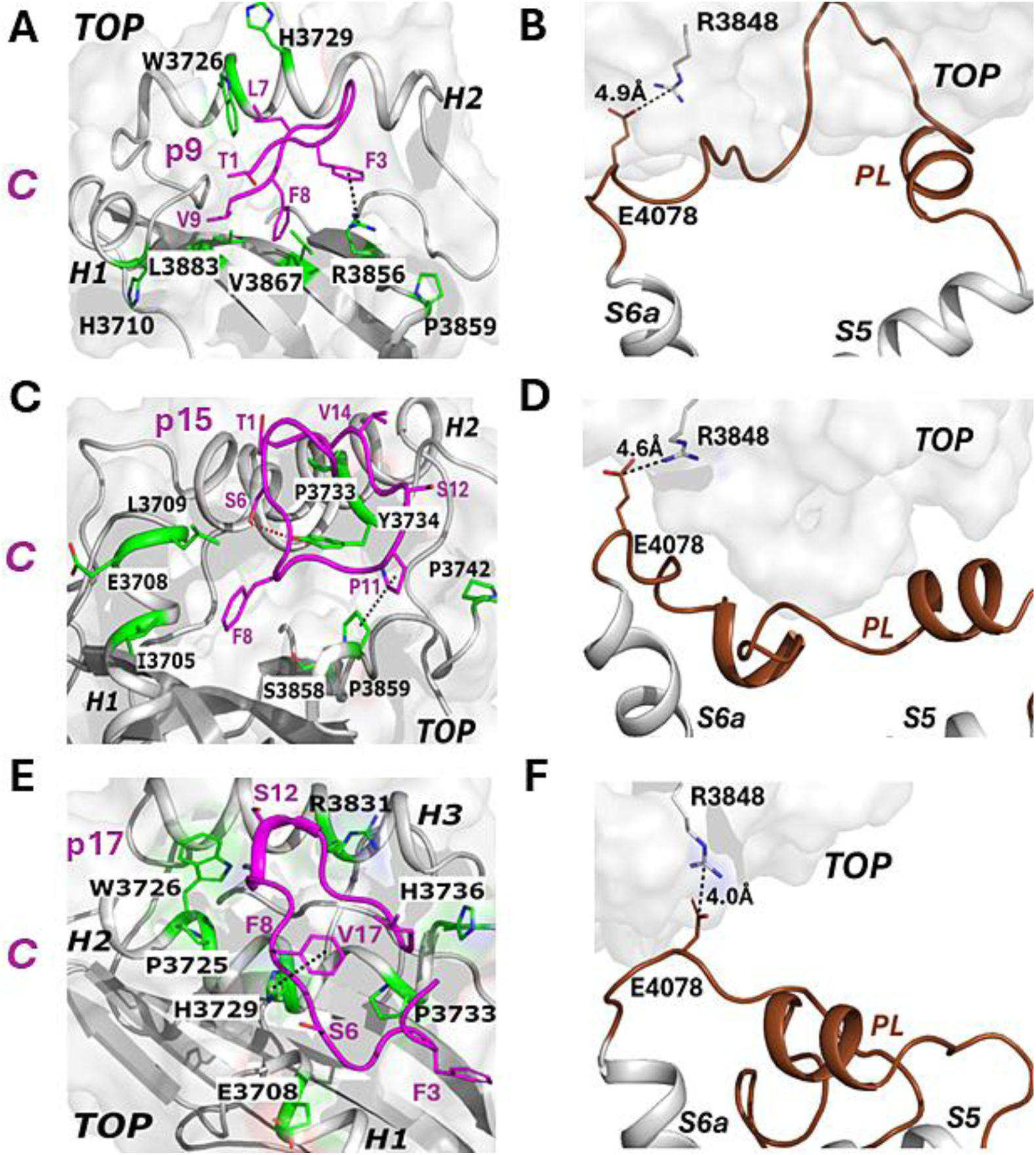
Representative low-energy conformational states of p9, p15 and p17 binding to CTF-R3063C: Interactions observed between peptide-protein residues in Closed (C) states of **(A)** p9-bound, **(C)** p15-bound and **(E)** p17-bound CTF-R3063C. Distance between the TOP domain residue R3848 and pore loop (PL) residue E4078 observed in Closed (C) states of **(B)** p9-bound, **(D)** p15-bound and **(F)** p17-bound CTF-R3063C.

For the p15-bound CTF-R3063C mutant, the calculated free energy profile showed two low-energy states - ‘I2’ and ‘Closed’ with an additional I1 state **(Figure 4B)**. In the **I1** state, characterized by the distance observed between the TOP domain residue R3848 and PL residue E4078 around ∼11 Å **(Figure S10D)**, p15 retained a linear conformation and remained bound between the *H1* and *H2* helices **(Figure S10C)** and formed two aromatic interactions between P10-H3736 and H13-H3736 peptide-protein residue pairs **(Figure S10C)**. In the **I2** state, the distance between the TOP domain residue R3848 and PL residue E4078 was centered around ∼25 Å **(Figure S10F)** while p15 retained a similar linear conformation as the Intermediate I1 state, however, the binding site was shifted to between the Finger-1 motif and the *H3* helix in the TOP domain **(Figure S10E)** where it formed one cation-π interaction between F3-R3833 and one aromatic interaction between H13-P3760 peptide-protein residue pairs **(Figure S10E)**. The **Closed** state was characterized by the formation of the salt bridge interaction between the TOP domain residue R3848 and PL residue E4078 with ∼4.6 Å distance **(Figure 5D)**. p15 also adopted a cyclized conformation with the T1 and V14 residues in close proximity and remained bound to the TOP domain primarily to the *H1* and *H2* region **(Figure 5C)**. In the Closed state, p15 formed one hydrogen-bonded interaction between S6-Y3734 along with one hydrophobic interaction between P11-P3859 peptide-protein pairs **(Figure 5C)**. For p15-bound CTF-R3063C mutant, we also observed the formation of the hydrogen-bonded interaction between the stalk residue N3074 and TM5 residue S3585 consistent with the closed/active conformation state **(Figure S7E and S8E)**.

Similarly, for the p17-bound CTF-R3063C mutant, the free energy profile also showed two low-energy states-‘I2’ and ‘Closed’ with an additional ‘Open’ state **(Figure 4C)**. In the **Open** state, the important signaling activation interaction between the TOP domain residue R3848 and PL residue E4078^50, 51^ remained separated from each other at ∼30 Å distance even though p17 remained bound to the extracellular TOP domain between the *H1*-*H2* **(Figure 4C)**. The **I2** state was characterized by ∼19 Å distance between the TOP domain residue R3848 and PL residue E4078 **(Figure S10H)** with p17 adopting a linear conformation and its binding site is shifted between the Finger-1 motif and the *H3* helix in the TOP domain **(Figure S10G)**. p17 also formed one hydrophobic interaction between P11-P3760 and one aromatic interaction between H13-H3768 peptide-protein pairs **(Figure S10G)**.

In the **Closed** state, the important signaling activation interaction between the TOP domain residue R3848 and PL residue E4078^50, 51^ was formed at ∼4.0 Å distance **(Figure 5F)** while p17 adopted a cyclized conformation with residues F3 and V17 in close proximity and remained primarily bound to the *H1*-*H2* region in the TOP domain **(Figure 5E)**. p17 formed one aromatic interaction between the F8-H3729 peptide-protein residue pair **(Figure 5E)**. For p17-bound CTF-R3063C mutant, the hydrogen-bonded residue-residue interaction between the stalk residue N3074 and TM5 residue S3585 was also observed, consistent with the closed/active conformational state of wild type PC1 CTF **(Figure S7F and S8F)**.

### 3.5 Mutant tethered stalk-TOP interactions influenced differential activation of PC1 CTF mutants by stalk-derived peptide agonists

To determine why signaling by the CTF-G3052R mutant could only be rescued by the p17 stalk-derived peptide while all the three stalk-derived peptides could rescue signaling by CTF-R3063C mutant, we examined the interactions of the tethered stalk with the extracellular regions of CTF in each of the peptide-bound mutant CTF systems.

Pep-GaMD simulations on the p9/p15-bound CTF-G3052R mutant captured the interaction of the tethered stalk with the TOP domain **(Figure S11A, S11B and Movie S1)**. As p9 sampled the **I1** state, the tethered stalk bound between *H3*-*H2* region and formed one polar interaction between its residue R3052 and residue E3828 in *H3* **(Figure S12A)**. In the **I2** state of p9, the mutant stalk moved closer to the binding site between *H1* and *H2* helices and formed more extensive polar interactions between residue R3052 and residues L3709, H3710, S3711 and H3729 from the TOP domain **(Figure S12B)**. Similarly, as p15 sampled the **I1** state, the tethered stalk formed one polar interaction between its residue R3052 and D3304 residue in the ECL1 loop **(Figure S13A)**. In the **I2** state of p15, the stalk bound between the TOP domain and ECL1 loop and formed two polar interactions between its residue R3052 and A3301 and D3304 residues in the ECL1 loop **(Figure S13B)**. However, for the p17-bound CTF-G3052R mutant, the tethered stalk mostly moved away from the TOP domain **(Figure S11C and Movie S2)**, leading to disruption of interactions between R3052 and TOP domain residues **(Figure S14)**.

For p9-bound CTF-R3063C mutant, as p9 sampled the **I2** state, the tethered stalk bound between the *H3*-*H2* region and formed two polar interactions between its mutant residue C3063 and residue Q3739 in the TOP domain and residue V3312 from ECL1 loop **(Figure S15A)**. In contrast, as p9 sampled the **Closed** state, the tethered stalk formed one polar interaction between residue C3063 and residue H3311 from ECL1 loop **(Figure S15B)**, however no interactions were formed between C3063 and TOP domain residues **(Figure S11D and Movie S3)**. Similarly, for the p15-bound CTF-R3063C mutant, as p15 sampled the **I2** state, the tethered stalk bound between the TOP domain and ECL1 loop and formed one polar interaction between its residue C3063 and residue A3306 from ECL1 loop **(Figure S16B)**. However, as p15 sampled the **I1** and the **Closed** state, the tethered stalk mostly moved away from the TOP domain **(Figure S16A and S16C)** leading to disruption of interactions between C3063 and the TOP domain residues **(Figure S11E and Figure S16)**.

Similarly, for p17-bound CTF-R3063C mutant, as p17 sampled the **I2** state, the tethered stalk bound between the TOP domain and ECL1 loop and formed two polar interactions between its residue C3063 and residues R3314 and L3315 from ECL1 loop **(Figure S17A)**. Finally, as p17 sampled the **Closed** state, the tethered stalk mostly moved away from the TOP domain, leading to disruption of interactions between C3063 and the TOP domain residues **(Figure S11F and Figure S17B)**. The above findings are consistent with the hypothesis that substitution of the small, nonpolar Gly with a polar Arg residue in the N-terminal region of the tethered stalk **(Figure 1A)** enabled the *de novo* formation of polar interactions that disrupted the allosteric activation pathway and prevented signaling activation by the shorter peptide agonists. This is in contrast to the substitution of Arg by Cys in the middle of the tethered stalk **(Figure 1A)** which was unable to maintain such spurious interactions in the presence of p9, p15 or p19 thus allowing CTF-R3063C re-activation by the peptide agonists.

The closed active state was observed for p17-bound CTF-G3052R as well as p9/p15/p17-bound CTF-R3063C mutants with the important signaling activation interaction^50, 51^ formed between the residues R3848 and E4078 in the TOP domain and PL. We also observed the formation of the hydrogen-bonded residue-residue interaction between N3074 residue from base of stalk and S3585 residue present in TM5 **(Figure S8C-F)** leading to a closed/active state with reduced stalk-TOP contacts **(Figure S18C-F)**. However, for the p9/p15-bound CTF-G3052R mutant systems where we observed the tethered stalk interacting with the TOP domain, the key residue interactions (R3848-E4078 and N3074-S3585) were observed to be in the open/inactive state with increased stalk-TOP contacts **(Figure S8A-B and S18A-B)**.

### 3.6 Analysis of *H1*-*H2* interhelical angle reveals further characteristics of agonist binding site

Previous studies implicated the *H1*–*H2* region on the TOP domain as the binding interface for the PC1 tethered stalk^50^ and the stalk-derived peptide agonists^51^, with additional contacts involving the β3, β4, and β5 strands^50^. Because both TA and stalk peptide binding occur between helices *H1* and *H2*, changes in the relative orientation of these helices may directly reflect structural rearrangements of the binding site. We compared the *in silico* closed active conformation of PC1 CTF^51^ with the cryo-EM structure (RCSB PDB: 6A70)^15^ and found differences in the interface between *H1*-*H2* in the TOP domain **(Figure S1B)**. In the active conformation of PC1 CTF^51^, the angle between *H1* and *H2* is ∼60 to 80 degrees **(Figure S19A)**, while in the cryo-EM structure^15^, which represents the inactive low free energy conformational state, the angle between *H1* and *H2* is 30 degrees **(Figure S19A).** The increase in the *H1-H2* interhelical angle from the inactive to the active conformation of PC1 CTF is suggestive of a less accessible/more restricted binding site which becomes more accessible/less restricted leading to the active low free energy conformational state. Therefore, the angle between the principal axes of *H1* and *H2* helices of the TOP domain was calculated to further characterize the peptide agonists’ binding site^50^.

Angle between the *H1*-*H2* helices for the p17-bound CTF-G3052R and p9/p15/p17-bound CTF-R3063C mutants were maintained around ∼60-80 degrees **(Figure S19D-G)** along with the formation of the active state **(Figure S20C-F)**. However, for p9/p15-bound CTF-G3052R mutant, the angle between the *H1*-*H2* helices at the peptide binding site was reduced to ∼20-40 degrees **(Figure S19B and S19C)** further characterizing the inactive state with its broken salt bridge interaction between R3848 and E4078 residues **(Figure S20A and S20B)**.

## 4. CONCLUSION

In ADPKD, 10 single nucleotide polymorphisms have been identified within the stalk of PC1 (ADPKD Variant Database; https://pkdb.mayo.edu/welcome), most of which were found to significantly reduce stalk TA-mediated activation of PC1 CTF to the NFAT reporter^64^. In particular, two ADPKD-associated missense variants, G3052R and R3063C, within the stalk region^51^ were previously shown to inhibit NFAT reporter activation in signaling assays and to be unable to sample the active conformational state in GaMD analyses^51^. Therefore, in this study, we chose these two mutant CTF proteins to investigate the ability of selective stalk-derived peptide agonists to rescue signaling and to reveal molecular binding and activation mechanisms.

We found p17 was able to re-activate signaling by both CTF-G3052R and -R3063C mutants, while p9 and p15 could only rescue CTF-R3063C signaling. To reveal the molecular binding mechanisms of the soluble stalk-derived peptides, we employed peptide docking using HPEPDOCK^56^ and Pep-GaMD^53^ simulations to examine CTF protein-peptide agonist interactions. Pep-GaMD simulations^53^ were able to refine the docking conformations of the three peptide agonists bound to the two CTF stalk mutants. Importantly, the simulation findings correlated with the experimental cellular signaling data, in that the low-energy active conformation was observed between p17 and both CTF-G3052R and -R3063C mutants and in p9-R3063C and p15-R3063C systems. The results of cell signaling and Pep-GaMD analyses are illustrated in **Figure 6**. In the p9/p15-bound CTF-G3052R mutant, the tethered stalk formed extensive TOP domain interactions, reducing the *H1*–*H2* angle (∼20–40°) and disrupting the R3848–E4078 and N3074–S3585 interactions, resulting in an inactive state **(Figure 6A)**. In contrast, the p17-bound CTF-G3052R and p9/p15/p17-bound CTF-R3063C mutants exhibited minimal stalk-TOP domain interactions, maintaining the *H1*–*H2* angle (∼60–80°) leading to the formation of the R3848–E4078 and N3074–S3585 interactions and corresponding to an active state **(Figure 6B, C)**. The peptide-binding interface was primarily stabilized by residues within the *H1*–*H2* region of the TOP domain **(Figure 6D)**.

**Figure 6.**
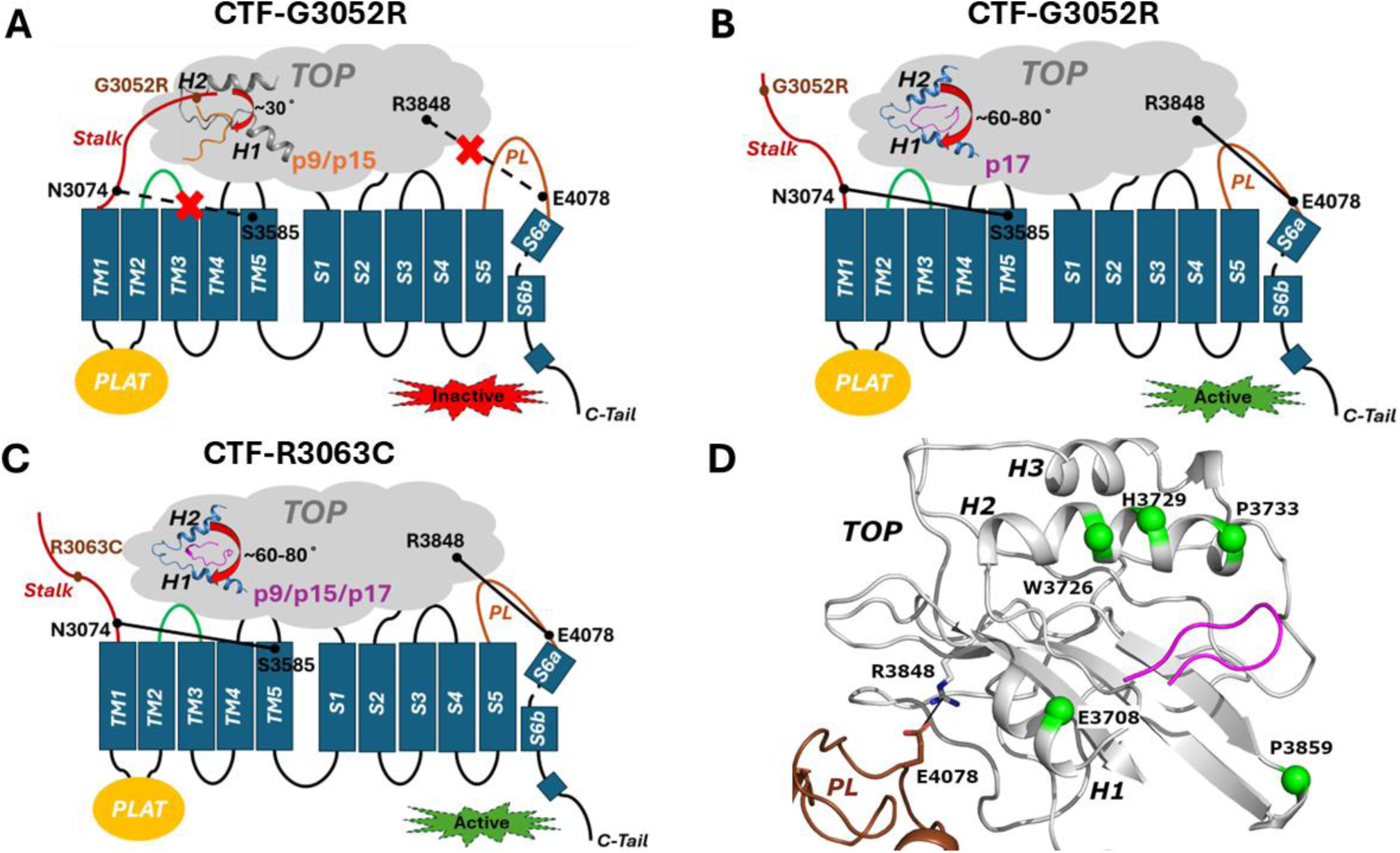
Models summarizing the salient features and resulting activation state of PC1 CTF disease-associated mutants with the stalk-derived peptide agonists: **(A)** p9/p15-bound CTF-G3052R mutant, **(B)** p17-bound CTF-G3052R mutant, **(C)** p9/p15/p17-bound CTF-R3063C mutant and **(D)** Structural model of the TOP domain highlighting key peptide-binding residues (W3726, E3708, H3729, P3733, and P3859) that contribute to stabilizing stalk-derived peptide agonists. Peptide binding alters the *H1*-*H2* angle and modulates the stalk-TOP-PL interactions (R3848–E4078 and N3074–S3585 interactions), to regulate the active and inactive states of PC1 CTF similar to the TA-mediated allosteric activation mechanism^51^.

Pep-GaMD simulations further provided mechanistic insights into how different stalk-derived peptide agonists regulate signaling in the ADPKD-associated PC1 CTF mutants G3052R and R3063C **(Figure 6)**. In the p9/p15-bound CTF-G3052R mutant, the peptide remained bound to the extracellular protein surface, while the stalk formed extensive interactions with the TOP domain. The angle between the *H1* and *H2* helices in the binding site reduced to ∼20-40 degrees, suggestive of a less accessible/more restricted binding site. Furthermore, the p9/p15-bound CTF-G3052R systems were characterized by the inactive low-energy conformational state that was represented by broken key residue interactions (R3848-E4078 and N3074-S3585) **(Figure 6A)**. In contrast, in simulations of p17-bound CTF-G3052R and p9/p15/p17-bound CTF-R3063C mutants, peptide binding primarily remained between the helices *H1* and *H2*, and the stalk moved away with minimal TOP domain contacts. The angle between the *H1*–*H2* was maintained at ∼60–80°, indicating a more accessible/less restricted binding site, and was characterized by the active low-energy conformational state with formation of the important R3848-E4078 and N3074-S3585 residue interactions **(Figure 6B and C**). These observations suggest that charged or polar missense mutations within the N-terminal region of the stalk may have a different and more significant impact on PC1 activation compared to mutating a residue in the middle (15^th^ position of 25 residue stalk) of the stalk sequence. Structural analysis further identified key TOP domain residues (W3726, H3729, P3733, E3708, and P3859) that contribute to the binding interface and stabilized the peptide binding **(Figure 6D)**. These key residues have also been shown to be critical in previous studies for binding of tethered stalk and stalk-derived peptides^50, 51^.

We observed similarities among the active conformational states of the stalk-derived peptide agonist-bound CTF-G3052R and CTF-R3063C mutants. Particularly, when inducing a closed/active CTF conformation, the three stalk-derived peptide agonists (p9, p15 and p15) acquired a circular conformation with terminal (or near terminal) residues coming into close proximity. Moreover, bulky residues of the peptides, i.e., F3, F8, P10, P11 and H13, inserted into hydrophobic grooves of the TOP domain surrounded by residues W3726, P3725, Y3734, P3859, Y3857 and P3733 located mainly in the *H2* helix and *β3*-*β4* turn elements. Furthermore, the peptides bound to the interface between the helices *H1*-*H2* on the TOP domain. Common interacting residues between CTF protein (E3708, W3726, H3729, P3733 and P3859) and peptide agonists (F3, F8, P10, S12 and V17) were identified for both G3052R and R3063C mutants. Key residues such as E3708, W3726 and P3859 have been previously reported, e.g., interactions of peptides p17 and p21 with E3708 were observed for the activation of stalkless CTF^50^. A Potts covariation analysis of residue interactions in PC1 CTF also revealed W3726 and P3859 in the TOP domain to be strongly interacting with stalk sequence^50^.

The binding interactions of the stalk derived peptides with PC1 CTF share similarities with those observed for the ADGRs. For ADGRs, their active conformation occurs when the tethered agonist inserts into the orthosteric site within the TM region and forms extensive hydrophobic interactions^43, 45, 46^. A hydrophobic F/Y/LXφφφXφ motif in the ADGR’s Stachel sequence forms five finger-like projections projecting into the hydrophobic pit of the TM bundle^43, 45^. For example, in ADGRG2-G_s_ complex, the lower rim of the Stachel peptide (F601, I603, L604, L605, D606 and L607) is inserted into the hydrophobic pocket and forms hydrophobic ring stacking interactions with F671^2.64^, L693^3.36^, Y779^5.36^, W838^6.53^, F842^6.57^, F856^7.42^ and W771^ECL2^ residues from the receptor’s binding pocket^45^. These findings are similar to our present study, in which the peptide agonists p9, p15 and p17 (F3, F8, P10, P11, S12, F16 and V17) bind to a hydrophobic region in the extracellular TOP domain of PC1 protein formed by residues P3725, W3726, H3729, H3736 and P3859 **(Table S7)**.

Altogether, this work demonstrates the potential for PC1 mutants with reduced signaling function to be rescued by stalk-derived peptide agonists as a novel therapeutic approach. Identification of the target peptide binding site in the TOP domain of PC1 CTF will allow the design of novel synthetic peptides or small molecules to aid in the rescue of PC1 residual function levels in the presence of ADPKD missense mutations. Based on the observation of compact circular conformations of bound stalk-derived peptides in the Pep-GaMD simulations, future studies will involve the design of macrocyclic peptides that are potentially more potent compared to linear peptides. Peptide design programs such as AFCycDesign^65^ and RFpeptides^66^ could be used for such an endeavor.

## Supporting information

The Supporting Information includes details about the GaMD and Pep-GaMD methodologies, Supplemental Tables S1 to S7, Supplemental Figures S1 to S20.

## 5. ASSOCIATED CONTENT

### Supporting Information

The Supporting Information is available free of charge at http://pubs.acs.org.

The Supporting Information includes details about the GaMD and Pep-GaMD methodologies, Supplemental Tables S1 to S7, Supplemental Figures S1 to S20 (PDF) and Supplemental Movies S1 to S3 (MP4).

## 6. AUTHOR INFORMATION

### Notes

The authors declare no competing financial interest.

## 7. ACKNOWLEDGMENTS

This work was supported in part by National Institutes of Health R01DK123590 (to R.L.M.) and R56DK135824 (to Y.M., R.L.M. and A.H.), as well as the startup project 27110 at University of North Carolina - Chapel Hill (to Y.M.). This work used supercomputing resources with allocation award TG-MCB180049 through the Advanced Cyberinfrastructure Coordination Ecosystem: Services & Support (ACCESS) program, which is supported by National Science Foundation grants #2138259, #2138286, #2138307, #2137603, and #2138296, and project M2874 through the National Energy Research Scientific Computing Center (NERSC), which is a U.S. Department of Energy Office of Science User Facility operated under Contract No. DE-AC02-05CH11231 and the Research Computing Cluster at the University of North Carolina at Chapel Hill (UNC-CH).

## REFERENCES

(1) Harris, P. C.; Torres, V. E. Genetic mechanisms and signaling pathways in autosomal dominant polycystic kidney disease. The Journal of clinical investigation 2014, 124 (6), 2315–2324.

(2) Torres, V. E.; Harris, P. C.; Pirson, Y. Autosomal dominant polycystic kidney disease. The Lancet 2007, 369 (9569), 1287–1301.

(3) Boletta, A.; Caplan, M. J. Physiologic mechanisms underlying polycystic kidney disease. Physiological reviews 2025, 105 (3), 1553–1607.

(4) Al-Orjani, Q.; Alshriem, L. A.; Gallagher, G.; Buqaileh, R.; Azizi, N.; AbouAlaiwi, W. Mechanistic insights into the pathogenesis of polycystic kidney disease. Cells 2025, 14 (15), 1203.

(5) Luo, L.; Roy, S.; Li, L.; Ma, M. Polycystic kidney disease: novel insights into polycystin function. Trends in molecular medicine 2023, 29 (4), 268–281.

(6) Maser, R. L.; Calvet, J. P.; Parnell, S. C. The GPCR properties of polycystin-1-A new paradigm. Frontiers in Molecular Biosciences 2022, 9, 1035507.

(7) Hardy, E. P.; Haider, A. N.; Patel, M. M.; Nesin, V.; Hoang, H. T.; Gostynska, S. E.; Berry, W. L.; Pioszak, A. A.; Ahmad, M.; Parnell, S. C. A heteromeric TRP channel that functions as a WNT-activated G protein-coupled receptor. Nature Communications 2026, 17 (1), 3233.

(8) Yang, Y.; Ehrlich, B. E. Structural studies of the C-terminal tail of polycystin-2 (PC2) reveal insights into the mechanisms used for the functional regulation of PC2. The Journal of Physiology 2016, 594 (15), 4141–4149.

(9) Miyakawa, A.; Ibarra, C.; Malmersjö, S.; Aperia, A.; Wiklund, P.; Uhlén, P. Intracellular calcium release modulates polycystin-2 trafficking. BMC nephrology 2013, 14 (1), 34.

(10) Qian, F.; Germino, F. J.; Cai, Y.; Zhang, X.; Somlo, S.; Germino, G. G. PKD1 interacts with PKD2 through a probable coiled-coil domain. Nature genetics 1997, 16 (2), 179–183.

(11) Tsiokas, L.; Kim, E.; Arnould, T.; Sukhatme, V. P.; Walz, G. Homo-and heterodimeric interactions between the gene products of PKD1 and PKD 2. Proceedings of the National Academy of Sciences 1997, 94 (13), 6965–6970.

(12) Yu, Y.; Ulbrich, M. H.; Li, M.-H.; Buraei, Z.; Chen, X.-Z.; Ong, A. C.; Tong, L.; Isacoff, E. Y.; Yang, J. Structural and molecular basis of the assembly of the TRPP2/PKD1 complex. Proceedings of the National Academy of Sciences 2009, 106 (28), 11558–11563.

(13) Nauli, S. M.; Alenghat, F. J.; Luo, Y.; Williams, E.; Vassilev, P.; Li, X.; Elia, A. E.; Lu, W.; Brown, E. M.; Quinn, S. J. Polycystins 1 and 2 mediate mechanosensation in the primary cilium of kidney cells. Nature genetics 2003, 33 (2), 129–137.

(14) Kim, S.; Nie, H.; Nesin, V.; Tran, U.; Outeda, P.; Bai, C.-X.; Keeling, J.; Maskey, D.; Watnick, T.; Wessely, O. The polycystin complex mediates Wnt/Ca2+ signalling. Nature cell biology 2016, 18 (7), 752–764.

(15) Su, Q.; Hu, F.; Ge, X.; Lei, J.; Yu, S.; Wang, T.; Zhou, Q.; Mei, C.; Shi, Y. Structure of the human PKD1-PKD2 complex. Science 2018, 361 (6406), eaat9819.

(16) Wang, Z.; Ng, C.; Liu, X.; Wang, Y.; Li, B.; Kashyap, P.; Chaudhry, H. A.; Castro, A.; Kalontar, E. M.; Ilyayev, L. The ion channel function of polycystin-1 in the polycystin-1/polycystin-2 complex. The EMBO Reports 2019, 20 (11), EMBR201948336.

(17) Torres, V. E.; Chapman, A. B.; Devuyst, O.; Gansevoort, R. T.; Grantham, J. J.; Higashihara, E.; Perrone, R. D.; Krasa, H. B.; Ouyang, J.; Czerwiec, F. S. Tolvaptan in patients with autosomal dominant polycystic kidney disease. New England Journal of Medicine 2012, 367 (25), 2407–2418.

(18) Torres, V. E.; Chapman, A. B.; Devuyst, O.; Gansevoort, R. T.; Perrone, R. D.; Koch, G.; Ouyang, J.; McQuade, R. D.; Blais, J. D.; Czerwiec, F. S. Tolvaptan in later-stage autosomal dominant polycystic kidney disease. New England Journal of Medicine 2017, 377 (20), 1930–1942.

(19) Patel, D. M.; Dahl, N. K. Long-term safety of tolvaptan in ADPKD: where do We Stand? Clinical Journal of the American Society of Nephrology 2021, 16 (1), 3–5.

(20) Ingelfinger, J. R. Tolvaptan and autosomal dominant polycystic kidney disease. Mass Medical Soc: 2017; Vol. 377, pp 1988–1989.

(21) Hopp, K.; Ward, C. J.; Hommerding, C. J.; Nasr, S. H.; Tuan, H.-F.; Gainullin, V. G.; Rossetti, S.; Torres, V. E.; Harris, P. C. Functional polycystin-1 dosage governs autosomal dominant polycystic kidney disease severity. The Journal of clinical investigation 2012, 122 (11), 4257–4273.

(22) Tan, Y.-C.; Blumenfeld, J.; Rennert, H. Autosomal dominant polycystic kidney disease: genetics, mutations and microRNAs. Biochimica et Biophysica Acta (BBA)-Molecular Basis of Disease 2011, 1812 (10), 1202–1212.

(23) Rossetti, S.; Kubly, V. J.; Consugar, M. B.; Hopp, K.; Roy, S.; Horsley, S. W.; Chauveau, D.; Rees, L.; Barratt, T. M.; Van’t Hoff, W. G. Incompletely penetrant PKD1 alleles suggest a role for gene dosage in cyst initiation in polycystic kidney disease. Kidney international 2009, 75 (8), 848–855.

(24) Cai, Y.; Fedeles, S. V.; Dong, K.; Anyatonwu, G.; Onoe, T.; Mitobe, M.; Gao, J.-D.; Okuhara, D.; Tian, X.; Gallagher, A.-R. Altered trafficking and stability of polycystins underlie polycystic kidney disease. The Journal of clinical investigation 2014, 124 (12), 5129–5144.

(25) Hofherr, A.; Wagner, C. J.; Watnick, T.; Köttgen, M. Targeted rescue of a polycystic kidney disease mutation by lysosomal inhibition. Kidney international 2016, 89 (4), 949–955.

(26) Krappitz, M.; Gallagher, A.-R.; Fedeles, S. Is it time to fold the cysts away? Trends in molecular medicine 2016, 22 (12), 997–999.

(27) Sandford, R.; Sgotto, B.; Aparicio, S.; Brenner, S.; Vaudin, M.; Wilson, R. K.; Chissoe, S.; Pepin, K.; Bateman, A.; Chothia, C. Comparative analysis of the polycystic kidney disease 1 (PKD1) gene reveals an integral membrane glycoprotein with multiple evolutionary conserved domains. Human molecular genetics 1997, 6 (9), 1483–1489.

(28) Nims, N.; Vassmer, D.; Maser, R. L. Transmembrane domain analysis of polycystin-1, the product of the polycystic kidney disease-1 (PKD1) gene: evidence for 11 membrane-spanning domains. Biochemistry 2003, 42 (44), 13035–13048.

(29) Parnell, S. C.; Magenheimer, B. S.; Maser, R. L.; Rankin, C. A.; Smine, A.; Okamoto, T.; Calvet, J. P. The Polycystic Kidney Disease-1 Protein, Polycystin-1, Binds and Activates Heterotrimeric G-Proteinsin Vitro. Biochemical and biophysical research communications 1998, 251 (2), 625–631.

(30) Parnell, S. C.; Magenheimer, B. S.; Maser, R. L.; Zien, C. A.; Frischauf, A.-M.; Calvet, J. P. Polycystin-1 activation of c-Jun N-terminal kinase and AP-1 is mediated by heterotrimeric G proteins. Journal of Biological Chemistry 2002, 277 (22), 19566–19572.

(31) Delmas, P.; Nomura, H.; Li, X.; Lakkis, M.; Luo, Y.; Segal, Y.; Fernández-Fernández, J. M.; Harris, P.; Frischauf, A.-M.; Brown, D. A. Constitutive activation of G-proteins by polycystin-1 is antagonized by polycystin-2. Journal of Biological Chemistry 2002, 277 (13), 11276–11283.

(32) Puri, S.; Magenheimer, B. S.; Maser, R. L.; Ryan, E. M.; Zien, C. A.; Walker, D. D.; Wallace, D. P.; Hempson, S. J.; Calvet, J. P. Polycystin-1 activates the calcineurin/NFAT (nuclear factor of activated T-cells) signaling pathway. Journal of Biological Chemistry 2004, 279 (53), 55455–55464.

(33) Hama, T.; Park, F. Heterotrimeric G protein signaling in polycystic kidney disease. Physiological genomics 2016, 48 (7), 429–445.

(34) Weston, B. S.; Bagnéris, C.; Price, R. G.; Stirling, J. L. The polycystin-1 C-type lectin domain binds carbohydrate in a calcium-dependent manner, and interacts with extracellular matrix proteins in vitro. Biochimica Et Biophysica Acta (BBA)-Molecular Basis of Disease 2001, 1536 (2-3), 161–176.

(35) Ibraghimov-Beskrovnaya, O.; Bukanov, N. O.; Donohue, L. C.; Dackowski, W. R.; Klinger, K. W.; Landes, G. M. Strong homophilic interactions of the Ig-like domains of polycystin-1, the protein product of an autosomal dominant polycystic kidney disease gene, PKD1. Human molecular genetics 2000, 9 (11), 1641–1649.

(36) Weston, B. S.; Malhas, A. N.; Price, R. G. Structure–function relationships of the extracellular domain of the autosomal dominant polycystic kidney disease-associated protein, polycystin-1. FEBS letters 2003, 538 (1-3), 8–13.

(37) Ha, K.; Nobuhara, M.; Wang, Q.; Walker, R. V.; Qian, F.; Schartner, C.; Cao, E.; Delling, M. The heteromeric PC-1/PC-2 polycystin complex is activated by the PC-1 N-terminus. Elife 2020, 9, e60684.

(38) Ponting, C.; Hofmann, K.; Bork, P. A latrophilin/CL-1-like GPS domain in polycystin-1. Current biology 1999, 9 (16), R585–R588.

(39) Qian, F.; Boletta, A.; Bhunia, A. K.; Xu, H.; Liu, L.; Ahrabi, A. K.; Watnick, T. J.; Zhou, F.; Germino, G. G. Cleavage of polycystin-1 requires the receptor for egg jelly domain and is disrupted by human autosomal-dominant polycystic kidney disease 1-associated mutations. Proceedings of the National Academy of Sciences 2002, 99 (26), 16981–16986.

(40) Kurbegovic, A.; Kim, H.; Xu, H.; Yu, S.; Cruanès, J.; Maser, R. L.; Boletta, A.; Trudel, M.; Qian, F. Novel functional complexity of polycystin-1 by GPS cleavage in vivo: role in polycystic kidney disease. Molecular and cellular biology 2014, 34 (17), 3341–3353.

(41) Maser, R. L.; Calvet, J. P. Adhesion GPCRs as a paradigm for understanding polycystin-1 G protein regulation. Cell. Signal. 2020, 72, 109637.

(42) Prömel, S.; Langenhan, T.; Araç, D. Matching structure with function: the GAIN domain of adhesion-GPCR and PKD1-like proteins. Trends in pharmacological sciences 2013, 34 (8), 470–478.

(43) Ping, Y.-Q.; Xiao, P.; Yang, F.; Zhao, R.-J.; Guo, S.-C.; Yan, X.; Wu, X.; Zhang, C.; Lu, Y.; Zhao, F. Structural basis for the tethered peptide activation of adhesion GPCRs. Nature 2022, 604 (7907), 763–770.

(44) Araç, D.; Boucard, A. A.; Bolliger, M. F.; Nguyen, J.; Soltis, S. M.; Südhof, T. C.; Brunger, A. T. A novel evolutionarily conserved domain of cell-adhesion GPCRs mediates autoproteolysis. The EMBO journal 2012, 31 (6), 1364–1378.

(45) Xiao, P.; Guo, S.; Wen, X.; He, Q.-T.; Lin, H.; Huang, S.-M.; Gou, L.; Zhang, C.; Yang, Z.; Zhong, Y.-N. Tethered peptide activation mechanism of the adhesion GPCRs ADGRG2 and ADGRG4. Nature 2022, 604 (7907), 771–778.

(46) Qu, X.; Qiu, N.; Wang, M.; Zhang, B.; Du, J.; Zhong, Z.; Xu, W.; Chu, X.; Ma, L.; Yi, C. Structural basis of tethered agonism of the adhesion GPCRs ADGRD1 and ADGRF1. Nature 2022, 604 (7907), 779–785.

(47) Barros-Álvarez, X.; Nwokonko, R. M.; Vizurraga, A.; Matzov, D.; He, F.; Papasergi-Scott, M. M.; Robertson, M. J.; Panova, O.; Yardeni, E. H.; Seven, A. B. The tethered peptide activation mechanism of adhesion GPCRs. Nature 2022, 604 (7907), 757–762.

(48) Stoveken, H. M.; Hajduczok, A. G.; Xu, L.; Tall, G. G. Adhesion G protein-coupled receptors are activated by exposure of a cryptic tethered agonist. Proceedings of the National Academy of Sciences 2015, 112 (19), 6194–6199.

(49) Joshi, K.; Miao, Y. Mechanism of Tethered Agonist Binding to an Adhesion G-Protein-Coupled Receptor. The Journal of Physical Chemistry B 2025, 130 (1), 56–67.

(50) Pawnikar, S.; Magenheimer, B. S.; Joshi, K.; Nevarez-Munoz, E.; Haldane, A.; Maser, R. L.; Miao, Y. Activation of polycystin-1 signaling by binding of stalk-derived peptide agonists. Elife 2024, 13, RP95992.

(51) Pawnikar, S.; Magenheimer, B. S.; Munoz, E. N.; Maser, R. L.; Miao, Y. Mechanism of tethered agonist-mediated signaling by polycystin-1. Proceedings of the National Academy of Sciences 2022, 119 (19), e2113786119.

(52) Wang, Y.; Wang, Z.; Pavel, M. A.; Ng, C.; Kashyap, P.; Li, B.; Morais, T. D.; Ulloa, G. A.; Yu, Y. The diverse effects of pathogenic point mutations on ion channel activity of a gain-of-function polycystin-2. Journal of Biological Chemistry 2023, 299 (5).

(53) Wang, J.; Miao, Y. Peptide Gaussian accelerated molecular dynamics (Pep-GaMD): Enhanced sampling and free energy and kinetics calculations of peptide binding. The Journal of chemical physics 2020, 153 (15).

(54) Miao, Y.; Feher, V. A.; McCammon, J. A. Gaussian accelerated molecular dynamics: unconstrained enhanced sampling and free energy calculation. Journal of chemical theory and computation 2015, 11 (8), 3584–3595.

(55) Miao, Y.; Sinko, W.; Pierce, L.; Bucher, D.; Walker, R. C.; McCammon, J. A. Improved reweighting of accelerated molecular dynamics simulations for free energy calculation. Journal of chemical theory and computation 2014, 10 (7), 2677–2689.

(56) Zhou, P.; Jin, B.; Li, H.; Huang, S.-Y. HPEPDOCK: a web server for blind peptide–protein docking based on a hierarchical algorithm. Nucleic acids research 2018, 46 (W1), W443–W450.

(57) Jo, S.; Kim, T.; Iyer, V. G.; Im, W. CHARMM-GUI: a web-based graphical user interface for CHARMM. Journal of computational chemistry 2008, 29 (11), 1859–1865.

(58) Huang, J.; Rauscher, S.; Nawrocki, G.; Ran, T.; Feig, M.; De Groot, B. L.; Grubmüller, H.; MacKerell Jr, A. D. CHARMM36m: an improved force field for folded and intrinsically disordered proteins. Nature methods 2017, 14 (1), 71–73.

(59) Vanommeslaeghe, K.; MacKerell Jr, A. CHARMM additive and polarizable force fields for biophysics and computer-aided drug design. Biochimica et Biophysica Acta (BBA)-General Subjects 2015, 1850 (5), 861–871.

(60) Darden, T.; York, D.; Pedersen, L. Particle mesh Ewald: An N log (N) method for Ewald sums in large systems. Journal of chemical physics 1993, 98 (12), 10089–10092.

(61) Roe, D. R.; Cheatham III, T. E. PTRAJ and CPPTRAJ: software for processing and analysis of molecular dynamics trajectory data. Journal of chemical theory and computation 2013, 9 (7), 3084–3095.

(62) Humphrey, W.; Dalke, A.; Schulten, K. VMD: visual molecular dynamics. Journal of molecular graphics 1996, 14 (1), 33–38.

(63) Chen, Y.; Hu, X.; Fan, W.; Shen, L.; Zhang, Z.; Liu, X.; Du, J.; Li, H.; Chen, Y.; Li, H. Fast density peak clustering for large scale data based on kNN. Knowledge-Based Systems 2020, 187, 104824.

(64) Magenheimer, B. S.; Munoz, E. N.; Ravichandran, J.; Maser, R. L. Constitutive signaling by the C-terminal fragment of polycystin-1 is mediated by a tethered peptide agonist. bioRxiv 2021, 2021.2008. 2020.457171.

(65) Rettie, S. A.; Campbell, K. V.; Bera, A. K.; Kang, A.; Kozlov, S.; Bueso, Y. F.; De La Cruz, J.; Ahlrichs, M.; Cheng, S.; Gerben, S. R. Cyclic peptide structure prediction and design using AlphaFold2. Nature Communications 2025, 16 (1), 4730.

(66) Rettie, S. A.; Juergens, D.; Adebomi, V.; Bueso, Y. F.; Zhao, Q.; Leveille, A. N.; Liu, A.; Bera, A. K.; Wilms, J. A.; Üffing, A. Accurate de novo design of high-affinity protein-binding macrocycles using deep learning. Nature Chemical Biology 2025, 21 (12), 1948–1956.

