## Supplementary material for "Activation of Disease-Associated Polycystin-1 Mutants by Binding of Stalk-derived Agonist Peptides": The Supporting Information includes details about the GaMD and Pep-GaMD methodologies, Supplemental Tables S1 to S7, Supplemental Figures S1 to S20.

### Table of Contents


**1.** Computational Methods

**1.1** Gaussian Accelerated Molecular Dynamics (GaMD)

**1.2** Peptide-Gaussian Accelerated Molecular Dynamics (Pep-GaMD)

**1.3** Energetic reweighting of Pep-GaMD simulations for free energy calculations

**Table S1.** Summary of the Pep-GaMD simulations performed on the CTF-G3052R in the presence of p9-bound stalk-derived peptide agonist.

**Table S2.** Summary of the Pep-GaMD simulations performed on the CTF-G3052R in the presence of p15-bound stalk-derived peptide agonist.

**Table S3.** Summary of the Pep-GaMD simulations performed on the CTF-G3052R in the presence of p17-bound stalk-derived peptide agonist.

**Table S4.** Summary of the Pep-GaMD simulations performed on the CTF-R3063C in the presence of p9-bound stalk-derived peptide agonist.

**Table S5.** Summary of the Pep-GaMD simulations performed on the CTF-R3063C in the presence of p15-bound stalk-derived peptide agonist.

**Table S6.** Summary of the Pep-GaMD simulations performed on the CTF-R3063C in the presence of p17-bound stalk-derived peptide agonist.

**Table S7.** Key interaction residues involved in stabilizing different low-energy states of peptide agonists (p9, p15 and p17) bound to CTF-G3052R and CTF-R3063C systems, respectively.

**Figure S1.** Conformations of the TOP Domain in PC1 CTF.

**Figure S2.** Peptide docking conformations and computational model for Pep-GaMD simulations of CTF-G3052R.

**Figure S3.** Peptide docking conformations and computational model for Pep-GaMD simulations of CTF-R3063C.

**Figure S4.** Time courses of the number of contacts between stalk-derived peptide agonists and extracellular domains of mutant CTF proteins calculated from Pep-GaMD simulations.

**Figure S5.** Time courses of the distance between the CZ atom of R3848 and the CD atom of E4078 of mutant CTF proteins calculated from Pep-GaMD simulations.

**Figure S6.** Expression verification of PC1 CTF stalk mutants.

**Figure S7.** Time courses of the distance between the CG atom of N3074 and the OG atom of S3585 in mutant CTF proteins calculated from Pep-GaMD simulations.

**Figure S8.** Two-dimensional free energy profiles regarding the N3074-S3585 distances versus the R3848-E4078 distances.

**Figure S9.** Representative low-energy conformational states of p15 binding to CTF-G3052R.

**Figure S10.** Representative low-energy conformational states of p9, p15 and p17 binding to CTF-R3063C.

**Figure S11.** Time courses of the number of contacts between the N-terminal region of Stalk (first 9 residues) and the TOP domain calculated from Pep-GaMD simulations.

**Figure S12.** Distortion of the TOP domain binding site in the presence of G3052R mutation in the p9-bound CTF-G3052R observed in Pep-GaMD simulations.

**Figure S13.** Distortion of the TOP domain binding site in the presence of G3052R mutation in the p15-bound CTF-G3052R observed in Pep-GaMD simulations.

**Figure S14.** TOP domain binding site maintained in Pep-GaMD simulations on p17-bound CTF-G3052R.

**Figure S15.** TOP domain binding site maintained in Pep-GaMD simulations on p9-bound CTF-R3063C.

**Figure S16.** TOP domain binding site maintained in Pep-GaMD simulations on p15-bound CTF-R3063C.

**Figure S17.** TOP domain binding site maintained in Pep-GaMD simulations on p17-bound CTF-R3063C.

**Figure S18.** Two-dimensional free energy profiles regarding the N3074-S3585 distances versus the Stalk-TOP contacts.

**Figure S19.** Time course plots for the angle between the *H1* and *H2* helices for CTF mutants with p9, p15, and p17.

**Figure S20.** Free energy profiles based on the angle between the *H1* and *H2* helices for CTF mutants with p9, p15, and p17.

**Movie S1**. Representative movie for p9-bound CTF-G3052R system.

**Movie S2**. Representative movie for p17-bound CTF-G3052R system.

**Movie S3**. Representative movie for p9-bound CTF-R3063C system.

#### 2. References

### 1. Computational Methods

#### 1.1 Gaussian Accelerated Molecular Dynamics (GaMD)

GaMD enhances the conformational sampling of biomolecules by adding a harmonic boost potential to smooth the system potential energy surface[1]. Consider a system with *N* atoms at positions $r \equiv \{\vec{r}_1, \cdots, \vec{r}_N\}$. When the system potential $V(\vec{r})$ is lower than the threshold energy *E*, a boost potential is added as:

$$\Delta V(\vec{r}) = \left(\frac{1}{2}k\big(E - V(\vec{r})\big)^2, V(\vec{r}) < E\right) \quad (1)$$

where $k$ is the harmonic force constant. The modified system potential, $V^*(\vec{r}) = V(\vec{r}) + \Delta V(\vec{r})$ is given by:

$$V^*(\vec{r}) = V(\vec{r}) + \frac{1}{2}k(E - V(\vec{r}))^2,\ V(\vec{r}) < E \quad (2)$$

Otherwise, when the system potential is above the threshold energy, i.e., $V(\vec{r}) \geq \mathrm{E}$, the boost potential is set to zero and $V^*(\vec{r}) = V(\vec{r})$.

In order to smooth the potential energy surface for enhanced sampling, the boost potential needs to satisfy the following criteria. First, for any two arbitrary potential values $V_1(\vec{r})$ and $V_2(\vec{r})$ found on the original energy surface, if $V_1(\vec{r}) < V_2(\vec{r})$, $\Delta V$ should be a monotonic function that does not change the relative order of the biased potential values, i.e., $V_1^*(\vec{r}) < V_2^*(\vec{r})$. By replacing $V^*(\vec{r})$ with Equation (2) and isolating *E*, we then obtain:

$$E < \frac{1}{2}[V_1(\vec{r}) + V_2(\vec{r})] + \frac{1}{k} \quad (3)$$

Second, if $V_1(\vec{r}) < V_2(\vec{r})$, the potential difference observed on the smoothened energy surface should be smaller than that of the original, i.e., $V_2^*(\vec{r}) - V_1^*(\vec{r}) < V_2(\vec{r}) - V_1(\vec{r})$. Similarly, by replacing $V^*(\vec{r})$ with Equation (2), we can derive:

$$E > \frac{1}{2}\,[V_1(\vec{r}) + V_2(\vec{r})] \qquad (4)$$

with $V_{min} \leq V_1(\vec{r}) < V_2(\vec{r}) \leq V_{max}$, we need to set the threshold energy $E$ in the following range by combining Equations (3) and (4):

$$V_{max} \leq E \leq V_{min} + \frac{1}{k} \qquad (5)$$

where $V_{min}$ and $V_{max}$ are the system minimum and maximum potential energies. To ensure that Equation (5) is valid, $V_{max} \leq V_{min} + \frac{1}{k}$ and $k$ has to satisfy:

$$k \leq \frac{1}{V_{max} - V_{min}} \qquad (6)$$

Let us define $k \equiv \frac{k_0}{V_{max} - V_{min}}$, then $0 < k_0 \leq 1$. $k_0$ determines the magnitude of the applied boost potential. With greater $k_0$, higher boost potential is added to the potential energy surface, which provides enhanced sampling of biomolecules across decreased energy barriers.

Third, the standard deviation of $\Delta V$ needs to be small enough (i.e., narrow distribution) to ensure accurate reweighting using cumulant expansion to the second order[2]:

$$\sigma_{\Delta V} = \sqrt{\left(\frac{\partial \Delta V}{\partial V}\Big|_{V=V_{avg}}\right)^2 \sigma_V^2} = k\left(E - V_{avg}\right)\sigma_V \leq \sigma_0, \qquad (7)$$

where $V_{avg}$ and $\sigma_V$ are the average and standard deviation of the system potential energies, $\sigma_{\Delta V}$ is the standard deviation of $\Delta V$ with $\sigma_0$ as a user-specified upper limit (e.g., $10k_B$T) for accurate reweighting. Provided Equation (5) that gives the range of threshold energy $E$, when $E$ is set to the lower bound $E = V_{max}$, we substitute in $E$ and $k$, and obtain:

$$k_0 \leq \left(\frac{\sigma_0}{\sigma_V} \cdot \frac{V_{max} - V_{min}}{V_{max} - V_{avg}}\right) \qquad (8)$$

Let us define the right-hand side in Equation (8) as:

$$k_0' = \frac{\sigma_0}{\sigma_V} \cdot \frac{V_{max} - V_{min}}{V_{max} - V_{avg}} \tag{9}$$

For efficient enhanced sampling with the highest possible acceleration, $k_0$ can then be set to its upper bound as:

$$k_0 = \min(1.0, k_0') = \min\left(1.0, \frac{\sigma_0}{\sigma_V} \cdot \frac{V_{max} - V_{min}}{V_{max} - V_{avg}}\right) \tag{10}$$

The larger $\sigma_{\Delta V}$ is obtained from the original potential energy surface (particularly for large biomolecules), the smaller $k_0$ may be applicable to allow for accurate reweighting. Alternatively, when the threshold energy *E* is set to its upper bound $E = V_{min} + \frac{1}{k}$ according to Equation (5), we substitute in *E* and *k* in Equation (7) and obtain:

$$k_0 \geq \left(1 - \frac{\sigma_0}{\sigma_V}\right) . \frac{V_{max} - V_{min}}{V_{avg} - V_{min}} \tag{11}$$

Let us define the right-hand side in Equation (11) as $k_0'' \equiv \left(1 - \frac{\sigma_0}{\sigma_V}\right) . \frac{V_{max} - V_{min}}{V_{avg} - V_{min}}$. Note that a smaller $k_0$ will give higher threshold energy *E*, but smaller force constant *k*. When $0 < k_0'' \leq 1$, $k_0$ can be set to either $k_0''$ for the highest threshold energy *E* or its upper bound 1.0 for the greatest force constant *k*. In this regard, $k_0 = k_0''$ is applied in the current GaMD. Otherwise, $k_0$ is calculated using Eqn. (11).

Given *E* and $k_0$, we can calculate the boost potential as:

$$\Delta V(\vec{r}) = \frac{1}{2} k_0 \frac{I}{V_{max} - V_{min}} (E - V(\vec{r}))^2, \quad V(\vec{r}) < E \tag{12}$$

Similar to aMD, GaMD provides options to add only the total potential boost $\Delta V_{\mathrm{p}}$, only dihedral potential boost $\Delta V_{\mathrm{D}}$, or the dual potential boost (both $\Delta V_{\mathrm{p}}$ and $\Delta V_{\mathrm{D}}$). The dual-boost simulation generally provides higher acceleration than the other two types of simulations for enhanced sampling[3, 4].

#### 1.2 Peptide-Gaussian Accelerated Molecular Dynamics (Pep-GaMD)

Based on GaMD[1, 5-8], "Selective" GaMD methodology such as Peptide GaMD (Pep-GaMD) was developed to efficiently explore the peptide binding and unbinding processes[9]. Consider a system of peptide *L* binding to a protein *P* in a biological environment *E*. The system comprises of *N* atoms with their coordinates $r \equiv \{\vec{r}_1, \cdots, \vec{r}_N\}$ and momenta $p \equiv \{\vec{p}_1, \cdots, \vec{p}_N\}$. The system Hamiltonian can be expressed as:

$$H(r,p) = K(p) + V(r) \tag{13}$$

where $K(p)$ and $V(r)$ are the system kinetic and total potential energies, respectively. Next, we decompose the potential energy into the following terms:

$$\begin{aligned} V(r) &= V_{P,b}(r_P) + V_{L,b}(r_L) + V_{E,b}(r_E) \\ &+ V_{PP,nb}(r_P) + V_{LL,nb}(r_L) + V_{EE,nb}(r_E) \\ &+ V_{PL,nb}(r_{PL}) + V_{PE,nb}(r_{PE}) + V_{LE,nb}(r_{LE}) \end{aligned} \tag{14}$$

where $V_{P,b}$, $V_{L,b}$ and $V_{E,b}$ are the bonded potential energies in protein *P*, peptide *L* and environment *E*, respectively. $V_{PP,nb}$, $V_{LL,nb}$ and $V_{EE,nb}$ are the self non-bonded potential energies in protein *P*, peptide $L$ and environment *E*, respectively. $V_{PL,nb}$, $V_{PE,nb}$ and $V_{LE,nb}$ are the non-bonded interaction energies between *P-L*, *P-E* and *L-E*, respectively. According to classical molecular mechanics force fields[10, 11], the non-bonded potential energies are usually calculated as:

$$V_{nb} = V_{elec} + V_{vdW}, \tag{15}$$

where $V_{elec}$ and $V_{vdW}$ are the system electrostatic and van der Waals potential energies. Presumably, peptide binding mainly involves both the bonded and non-bonded interaction energies of the peptide since peptides often undergo large conformational changes during binding to the target proteins. Thus, the peptide essential potential energy is $V_L(r) = V_{LL,b}(r_L) + V_{LL,nb}(r_L) + V_{PL,nb}(r_{PL}) + V_{LE,nb}(r_{LE})$.

In selective Pep-GaMD, we add a boost potential selectively to the peptide essential potential energy as:

$$\Delta V_L(r) = \begin{cases} \frac{1}{2} k_L \big(E_L - V_L(r)\big)^2, \ V_L(r) < E_L \\ 0, \ V_L(r) \geq E_L \end{cases} \tag{16}$$

where $E_L$ is the threshold energy for applying boost potential and $k_L$ is the harmonic constant. Pep-GaMD simulation parameters are derived similarly as in the GaMD algorithm. When $E$ is set to the lower bound as the system maximum potential energy ($E=V_{max}$), the effective harmonic force constant $k_0$ can be calculated as:

$$k_0 = \min(1.0, k_0') = \min(1.0, \frac{\sigma_0}{\sigma_V}\frac{V_{max}-V_{min}}{V_{max}-V_{avg}}), \tag{17}$$

where $V_{max}$, $V_{min}$, $V_{avg}$ and $\sigma_V$ are the maximum, minimum, average and standard deviation of the boosted system potential energy, and $\sigma_0$ is the user-specified upper limit of the standard deviation of $\Delta V$ (e.g., $10k_B$T) for proper reweighting. The harmonic constant is calculated as $k = k_0 \cdot \frac{1}{V_{max}-V_{min}}$ with $0 < k_0 \leq 1$ . Alternatively, when the threshold energy $E$ is set to its upper bound $E = V_{min} + \frac{1}{k}$, $k_0$ is set to:

$$k_0 = k_0'' \equiv (1 - \frac{\sigma_0}{\sigma_V}) \frac{V_{max}-V_{min}}{V_{avg}-V_{min}}, \tag{18}$$

if $k_0''$ is found to be between *0* and *1*. Otherwise, $k_0$ is calculated using Equation (18).

In addition to selectively boosting the peptide, another boost potential in Pep-GaMD is applied on the protein and solvent to enhance conformational sampling of the protein and facilitate peptide binding. The second boost potential is thus calculated using the total system potential energy other than the peptide potential energy as:

$$\Delta V_D(r) = \begin{cases} \frac{1}{2} k_D \left(E_D - V_D(r)\right)^2, & V_D(r) < E_D \\ 0, & V_D(r) \geq E_D \end{cases} \quad (19)$$

where $V_D$ is the total system potential energy other than the peptide essential potential energy, $E_D$ is the corresponding threshold energy for applying the second boost potential and $k_D$ is the harmonic constant. This leads to dual-boost selective Pep-GaMD with the total boost potential $\Delta V(r) = \Delta V_L(r) + \Delta V_D(r)$.

##### 1.3 Energetic reweighting of Pep-GaMD simulations for free energy calculations

To perform energetic reweighting of Pep-GaMD simulations[9], the probability distribution defined along a selected reaction coordinate can be denoted as $p^*(A)$. Given $\Delta V(r)$ as the boost potential for each frame in Pep-GaMD simulations[9], $p^*(A)$ can be reweighted to recover the canonical ensemble distribution, $p(A)$, as follows:

$$p(A_j) = p^*(A_j) \frac{\langle e^{\beta \Delta V(\vec{r})} \rangle_j}{\sum_{i=1}^{M} \langle p^*(A_i) e^{\beta \Delta V(\vec{r})} \rangle_i}, \ j = 1, \ldots, M \quad (20)$$

where $M$ denotes the number of bins, $\beta = k_B T$ and $\langle e^{\beta \Delta V(\vec{r})} \rangle_j$ denotes the ensemble-averaged Boltzmann factor of $\Delta V(\vec{r})$ for the simulation frames observed in the $j^{\text{th}}$ bin. In order to reduce the energetic noise, the ensemble-averaged reweighting factor can be approximated using the cumulant expansion[12]:

$$\langle e^{\beta \Delta V(\vec{r})} \rangle = exp\left\{ \sum_{k=1}^{\infty} \frac{\beta^k}{k!} C_k \right\} \quad (21)$$

where the first two cumulants are defined as:

$$C_1 = \langle \Delta V \rangle \quad (22)$$

$$C_2 = \langle \Delta V^2 \rangle - \langle \Delta V \rangle^2 = \sigma_{\Delta V}^2$$

The boost potential obtained from Pep-GaMD simulations[9] usually shows near-Gaussian distribution. Therefore, cumulant expansion to the second order[12] provides a more accurate reweighting. The reweighted free energy $F(A) = -k_B T \ln p(A)$ is denoted as:

$$F(A) = F^*(A) - \sum_{k=1}^{2} \frac{\beta^k}{k!} C_k + F_c \tag{23}$$

where $F^*(A) = -k_B T \ln p^*(A)$ denotes the modified free energy obtained from Pep-GaMD simulation and $F_c$ is a constant.

**Table S1. Summary of the Pep-GaMD simulations performed on the CTF-G3052R in the presence of p9-bound stalk-derived peptide agonist.**

| System | System Size | ID | Simulation length | Boost Potential (kcal/mol) |
|---|---|---|---|---|
| **p9-CTF-G3052R** | 228,276 atoms | Sim1 | 1000ns | 13.26+/-4.06 |
| | | Sim2 | | 12.27+/-4.03 |
| | | Sim3 | | 12.95+/-4.02 |
| | | Sim4 | | 12.14+/-3.99 |
| | | Sim5 | | 13.64+/-4.10 |

**Table S2. Summary of the Pep-GaMD simulations performed on the CTF-G3052R in the presence of p15-bound stalk-derived peptide agonist.**

| System | System Size | ID | Simulation length | Boost Potential (kcal/mol) |
|---|---|---|---|---|
| **p15-CTF-G3052R** | 232,145 atoms | Sim1 | 1000ns | 11.88 ± 3.97 |
| | | Sim2 | | 12.44 ± 4.05 |
| | | Sim3 | | 11.61 ± 3.97 |
| | | Sim4 | | 12.28 ± 3.97 |
| | | Sim5 | | 12.28 ± 3.98 |

**Table S3. Summary of the Pep-GaMD simulations performed on the CTF-G3052R in the presence of p17-bound stalk-derived peptide agonist.**

| System | System Size | ID | Simulation length | Boost Potential (kcal/mol) |
|---|---|---|---|---|
| **p17-CTF-G3052R** | 211,144 atoms | Sim1 | 1000ns | 12.84 ± 4.01 |
| | | Sim2 | | 13.07 ± 4.02 |
| | | Sim3 | | 11.87 ± 3.97 |
| | | Sim4 | | 11.66 ± 3.93 |
| | | Sim5 | | 12.42 ± 4.00 |

**Table S4. Summary of the Pep-GaMD simulations performed on the CTF-R3063C in the presence of p9-bound stalk-derived peptide agonist.**

| System | System Size | ID | Simulation length | Boost Potential (kcal/mol) |
|---|---|---|---|---|
| **p9-CTF-R3063C** | 226,170 atoms | Sim1 | 1000ns | 9.95 ± 3.81 |
| | | Sim2 | | 11.26 ± 3.86 |
| | | Sim3 | | 11.07 ± 3.86 |
| | | Sim4 | | 11.86 ± 4.09 |
| | | Sim5 | | 11.35 ± 3.88 |

**Table S5. Summary of the Pep-GaMD simulations performed on the CTF-R3063C in the presence of p15-bound stalk-derived peptide agonist.**

| System | System Size | ID | Simulation length | Boost Potential (kcal/mol) |
|---|---|---|---|---|
| **p15-CTF-R3063C** | 228,881 atoms | Sim1 | 1000ns | 11.23 ± 3.87 |
| | | Sim2 | | 11.54 ± 4.00 |
| | | Sim3 | | 11.59 ± 3.95 |
| | | Sim4 | | 11.14 ± 3.89 |
| | | Sim5 | | 11.63 ± 4.02 |

**Table S6. Summary of the Pep-GaMD simulations performed on the CTF-R3063C in the presence of p17-bound stalk-derived peptide agonist.**

| System | System Size | ID | Simulation length | Boost Potential (kcal/mol) |
|---|---|---|---|---|
| **p17-CTF-R3063C** | 231,794 atoms | Sim1 | 1000ns | 12.60 ± 4.02 |
| | | Sim2 | | 12.81 ± 4.06 |
| | | Sim3 | | 12.57 ± 4.01 |
| | | Sim4 | | 13.03 ± 4.06 |
| | | Sim5 | | 12.67 ± 4.16 |

**Table S7. Key interaction residues involved in stabilizing different low-energy states of peptide agonists (p9, p15 and p17) bound to CTF-G3052R and CTF-R3063C systems, respectively.**

| System | I1 | | I2 | | Closed | |
|---|---|---|---|---|---|---|
| | Peptide | PC1 CTF | Peptide | PC1 CTF | Peptide | PC1 CTF |
| p9-bound CTF-G3052R | T1, S6, L7, F8 and V9 | K3706$^{H1}$, H3710$^{H1}$, P3725$^{H2}$, H3729$^{H2}$, and P3733$^{H2}$ | T1, A5, S6 and V9 | Q3702$^{H1}$, K3706$^{H1}$, H3729$^{H2}$, P3733$^{H2}$ and R3856$^{\beta3}$ | - | - |
| p15-bound CTF-G3052R | A2, F3, A5, L7 and H13 | A3306$^{ECL1}$, Y3307$^{ECL1}$ and S3308$^{ECL1}$ | T1, A2, A5, S6, L7 and F8 | L3701$^{H1}$, Q3702$^{H1}$, I3705$^{H1}$, K3706$^{H1}$ and L3709$^{H1}$ | - | - |
| p17-bound CTF-G3052R | F3, L7, P11, H13, R15 and F16 | W3724$^{H2}$, P3725$^{H2}$, W3726$^{H2}$, H3729$^{H2}$, P3733$^{H2}$, H3736$^{H2}$ and R3831$^{H3}$ | - | - | F3, F8, P10, S12, H13 and V17 | **E3708$^{H1}$**, R3712$^{Luminal\text{-}Loop}$, E3722$^{H2}$, P3725$^{H2}$, **W3726$^{H2}$**, H3729$^{H2}$, and H3736$^{H2}$ |
| p9-bound CTF-R3063C | - | - | T1, F3, A5, L7, F8 and V9 | Y3699$^{H1}$, R3700$^{H1}$, S3703$^{H1}$, K3706$^{H1}$, H3710$^{H1}$, F3714$^{Luminal\text{-}Loop}$, P3725$^{H2}$, W3726$^{H2}$, H3729$^{H2}$ and R3856$^{\beta3}$ | T1, F3, L7, F8 and V9 | H3710$^{H1}$, **W3726$^{H2}$**, H3729$^{H2}$, R3856$^{\beta3}$, **P3859**$^{\beta3\text{-}\ \beta4\text{-Turn}}$, V3867$^{\beta4}$ and L3883$^{\beta5}$ |
| p15-bound CTF-R3063C | S6, L7, F8, P10, P11, and H13 | Q3702$^{H1}$, E3708$^{H1}$, Y3734$^{H2}$, H3736$^{H2\text{-}\beta1\text{-Turn}}$, S3740$^{H2\text{-}\beta1\text{-Turn}}$, S3741$^{H2\text{-}\beta1\text{-Turn}}$, P3859$^{\beta3\text{-}\beta4\text{-Turn}}$ and V3861$^{\beta3\text{-}\beta4\text{-Turn}}$ | F3, L7, V9, P10, P11, S12, H13 and V14 | R3753$^{Finger\text{-}1}$, Q3755$^{Finger\text{-}1}$, Y3759$^{Finger\text{-}1}$, P3760$^{Finger\text{-}1}$, Y3819$^{Finger\text{-}1}$ and R3833$^{H3}$ | T1, S6, F8, P11, S12 and V14 | I3705$^{H1}$, **E3708$^{H1}$**, L3709$^{H1}$, P3733$^{H2}$, Y3734$^{H2}$, P3742$^{H2\text{-}\beta1\text{-Turn}}$, S3858$^{\beta3\text{-}\beta4\text{-Turn}}$ and **P3859**$^{\beta3\text{-}\beta4\text{-Turn}}$ |
| p17-bound CTF-R3063C | - | - | A2, A5, S6, F8, P11 and H13 | R3748$^{Finger\text{-}1}$, P3760$^{Finger\text{-}1}$, P3762$^{Finger\text{-}1}$, H3768$^{Finger\text{-}1}$, E3786$^{Finger\text{-}1}$, H3840$^{H3}$ and S3847$^{H3\text{-}\beta3\text{-Turn}}$ | F3, S6, F8, S12 and V17 | **E3708$^{H1}$**, P3725$^{H2}$, **W3726$^{H2}$**, H3729$^{H2}$, P3733$^{H2}$, H3736$^{H2}$ and R3831$^{H3}$ |

**Key Interacting Residues:** E3708$^{H1}$, W3726$^{H2}$ and P3859$^{\beta3\text{-}\beta4\text{-Turn}}$

**Figure S1. Conformations of the TOP Domain in PC1 CTF: (A)** Structural features of the PC1 TOP domain modeled from PC2 TOP domain[13]. **(B)** Comparison of the TOP domain conformations upon stalk binding in active CTF (shown as blue cartoon) with stalk[14] versus cryo-EM CTF (PDB: 6A70)[15] (shown as grey cartoon).

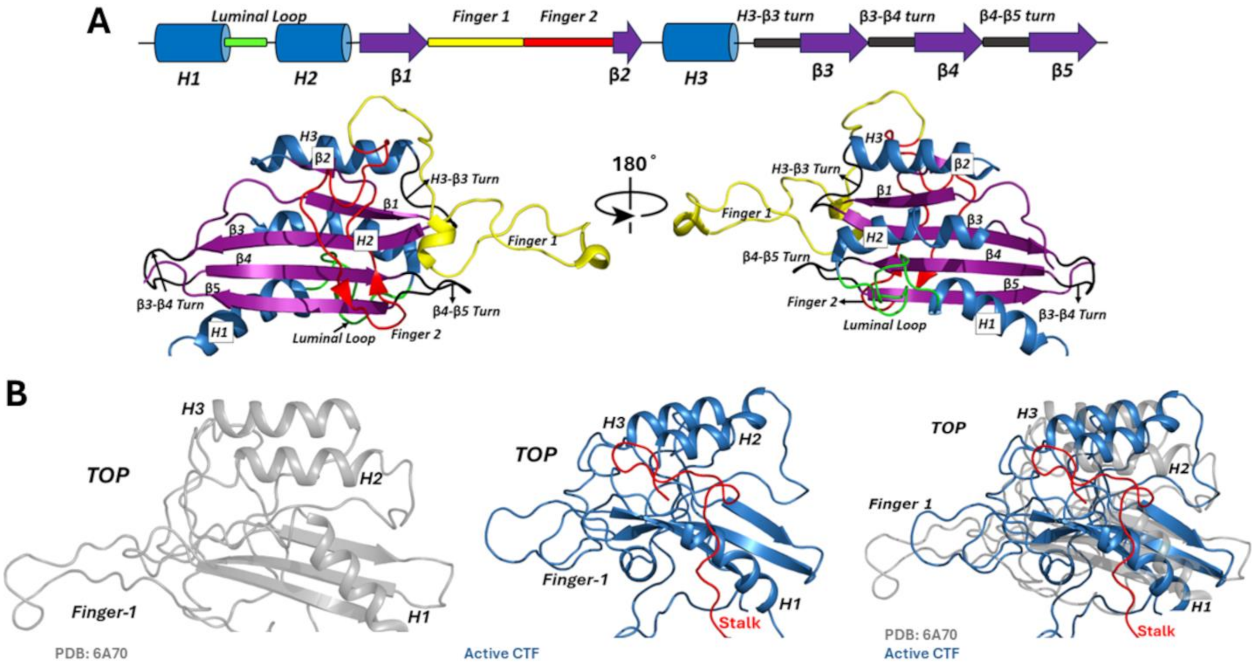

**Figure S2. Peptide docking conformations and computational model for Pep-GaMD simulations of CTF-G3052R: (A–B)** Docking conformations of p9, p15 and p17 to CTF-G3052R. **(C)** Simulation system of CTF-G3052R (grey cartoon) embedded in a palmitoyl-oleoyl-phosphatidyl-choline (POPC) lipid bilayer (cyan sticks) solvated in 0.15 M NaCl solution.

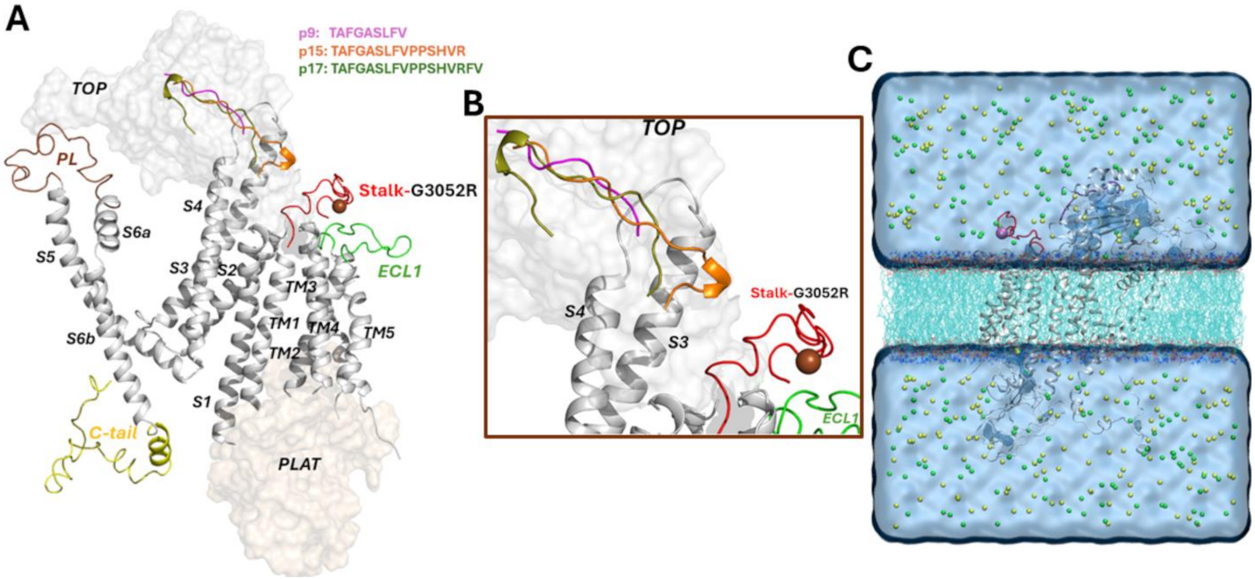

**Figure S3. Peptide docking conformations and computational model for Pep-GaMD simulations of CTF-R3063C: (A–B)** Docking conformations of p9, p15 and p17 to CTF-R3063C. **(C)** Simulation system of CTF-R3063C (grey cartoon) embedded in a palmitoyl-oleoyl-phosphatidyl-choline (POPC) lipid bilayer (cyan sticks) solvated in 0.15 M NaCl solution.

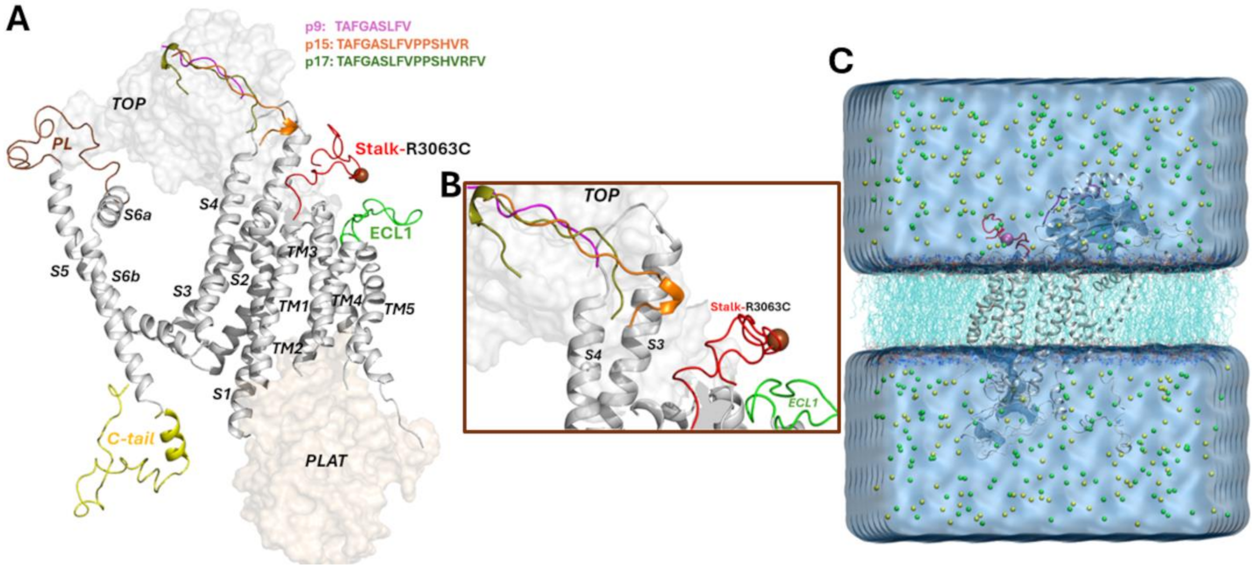

**Figure S4. Time courses of the number of contacts between stalk-derived peptide agonists and extracellular domains of mutant CTF proteins calculated from Pep-GaMD simulations of the: (A)** p9-bound CTF-G3052R, **(B)** p15-bound CTF-G3052R, **(C)** p17-bound CTF-G3052R, **(D)** p9-bound CTF-R3063C, **(E)** p15-bound CTF-R3063C, and **(F)** p17-bound CTF-R3063C systems, respectively. Residue contacts were calculated for heavy atoms with a distance cutoff of 4 Å.

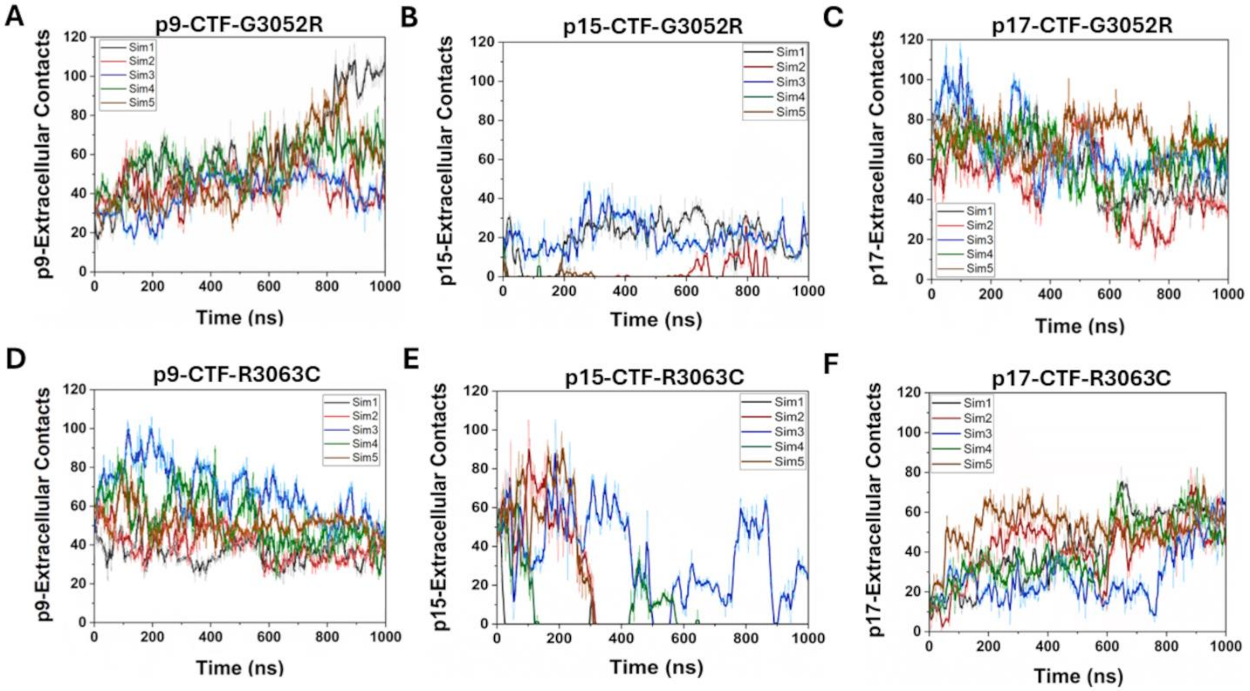

**Figure S5. Time courses of the distance between the CZ atom of R3848 and the CD atom of E4078 of mutant CTF proteins calculated from Pep-GaMD simulations of the: (A)** p9-bound CTF-G3052R, **(B)** p15-bound CTF-G3052R, **(C)** p17-bound CTF-G3052R, **(D)** p9-bound CTF-R3063C, **(E)** p15-bound CTF-R3063C, and **(F)** p17-bound CTF-R3063C systems, respectively.

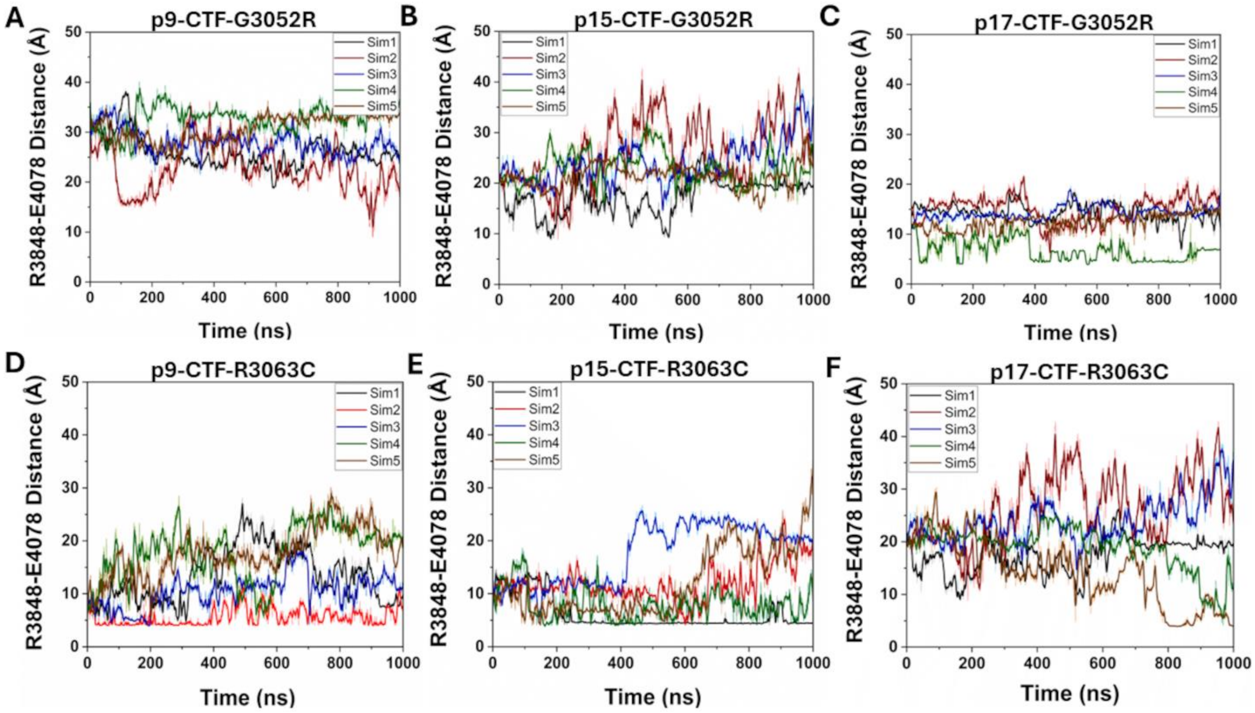

**Figure S6. Expression verification of PC1 CTF stalk mutants: (A)** Representative Western blot analysis of lysates from cells transfected with empty expression vector (ev), Δstalk CTF (Δst) or CTF-G3052R either as control (con, medium change only), or treated with p9, p15 or p17. **(B-C)** Representative Western blot analyses of lysates from cells transfected with empty expression vector (ev), Δstalk CTF (Δst) or CTF-R3063C either as control (con; medium change only), or treated with p9, p15 or p17.

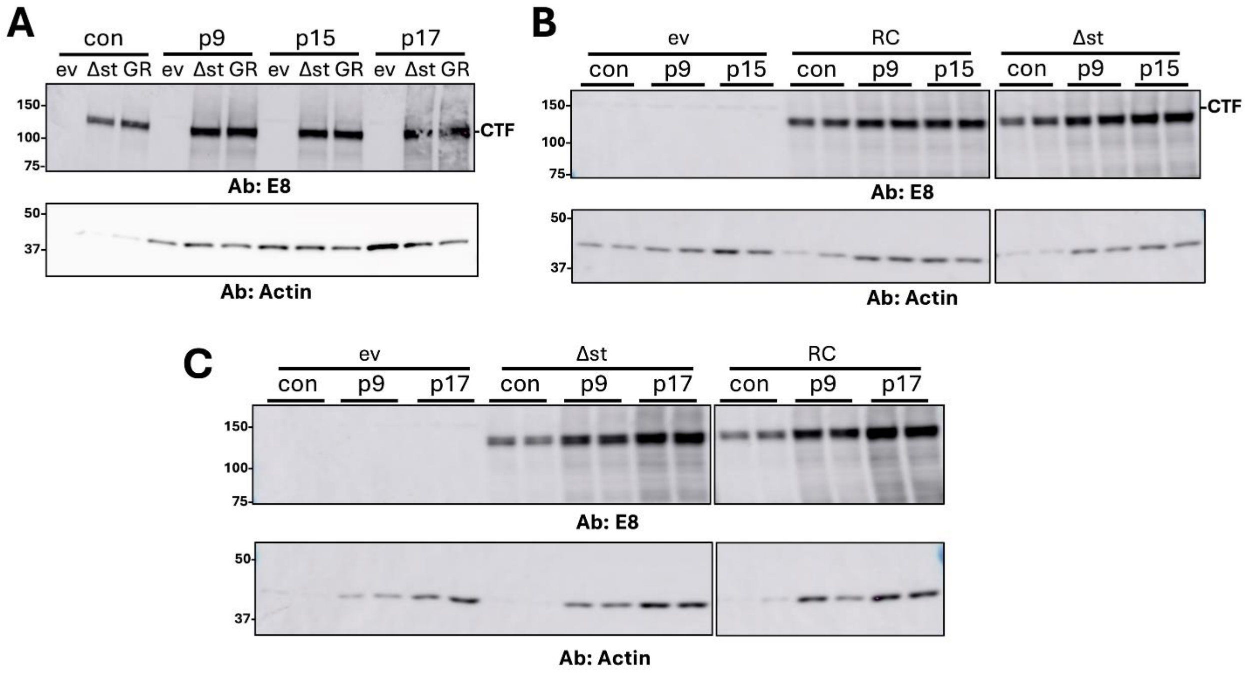

**Figure S7. Time courses of the distance between the CG atom of N3074 and the OG atom of S3585 in mutant CTF proteins calculated from Pep-GaMD simulations of the: (A)** p9-bound CTF-G3052R, **(B)** p15-bound CTF-G3052R, **(C)** p17-bound CTF-G3052R, **(D)** p9-bound CTF-R3063C, **(E)** p15-bound CTF-R3063C, and **(F)** p17-bound CTF-R3063C systems, respectively.

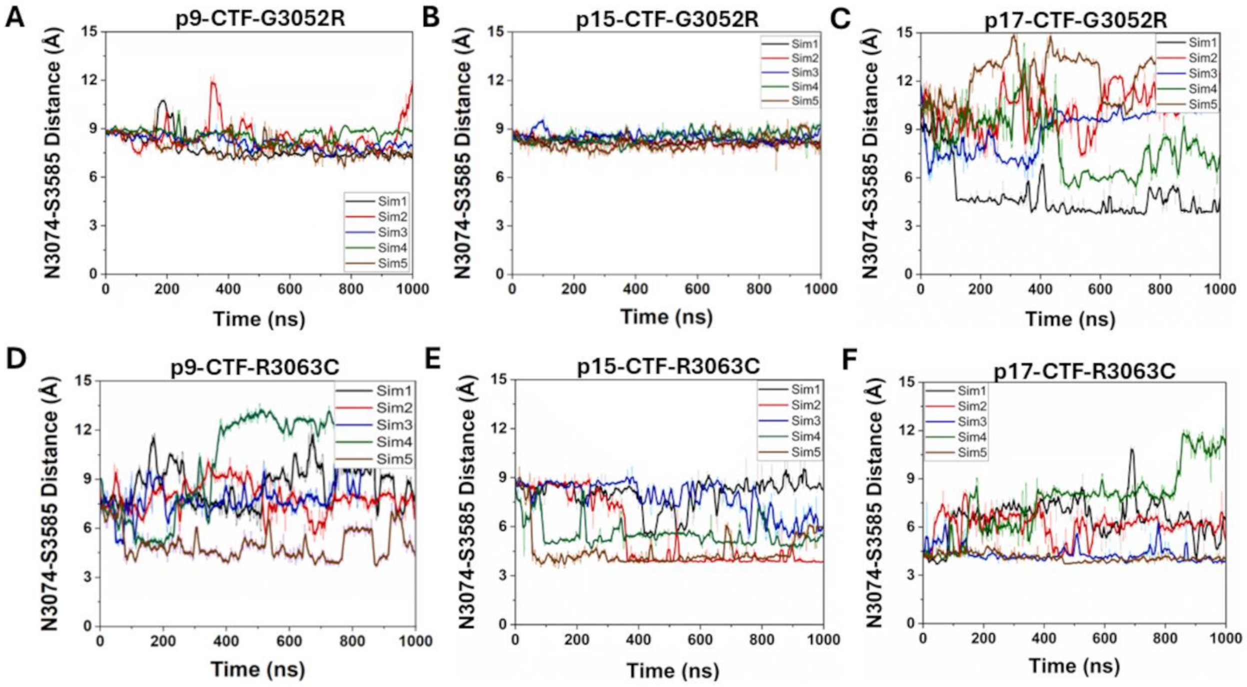

**Figure S8. Two-dimensional free energy profiles regarding the N3074-S3585 distances versus the R3848-E4078 distances of the: (A)** p9-bound CTF-G3052R, **(B)** p15-bound CTF-G3052R, **(C)** p17-bound CTF-G3052R, **(D)** p9-bound CTF-R3063C, **(E)** p15-bound CTF-R3063C, and **(F)** p17-bound CTF-R3063C systems. The distances between the CG atom in N3074 and the OG atom in S3585 and between the CZ atom in R3848 and the CD atom in E4078 were calculated from the Pep-GaMD simulations. Important low-energy conformational states are identified, including the "Open", "Intermediate I1," "Intermediate I2," and "Closed".

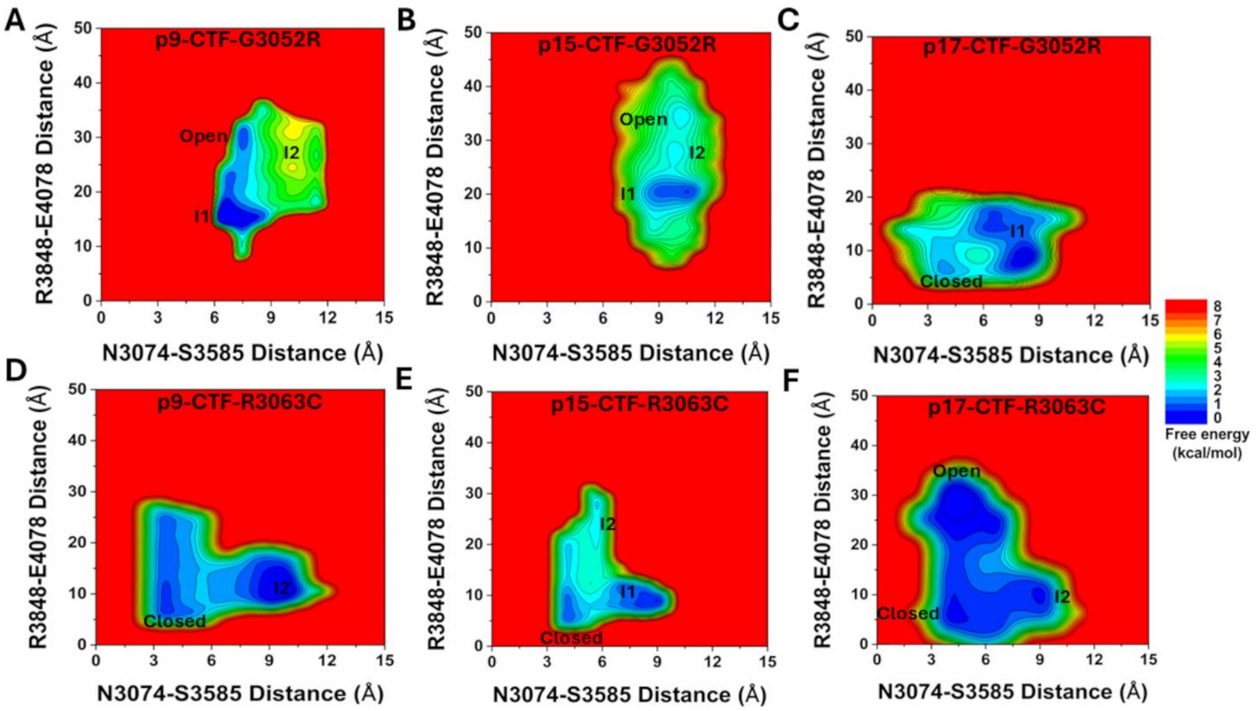

**Figure S9. Representative low-energy conformational states of p15 binding to CTF-G3052R: (A)** Interactions observed between peptide-protein residues and **(B)** the distance between the TOP domain residue R3848 and pore loop (PL) residue E4078 observed in Intermediate 1 (I1) state of p15-bound CTF-G3052R. **(C)** Interactions observed between peptide-protein residues and **(D)** the distance between the TOP domain residue R3848 and pore loop (PL) residue E4078 observed in Intermediate 2 (I2) state of p15-bound CTF-G3052R.

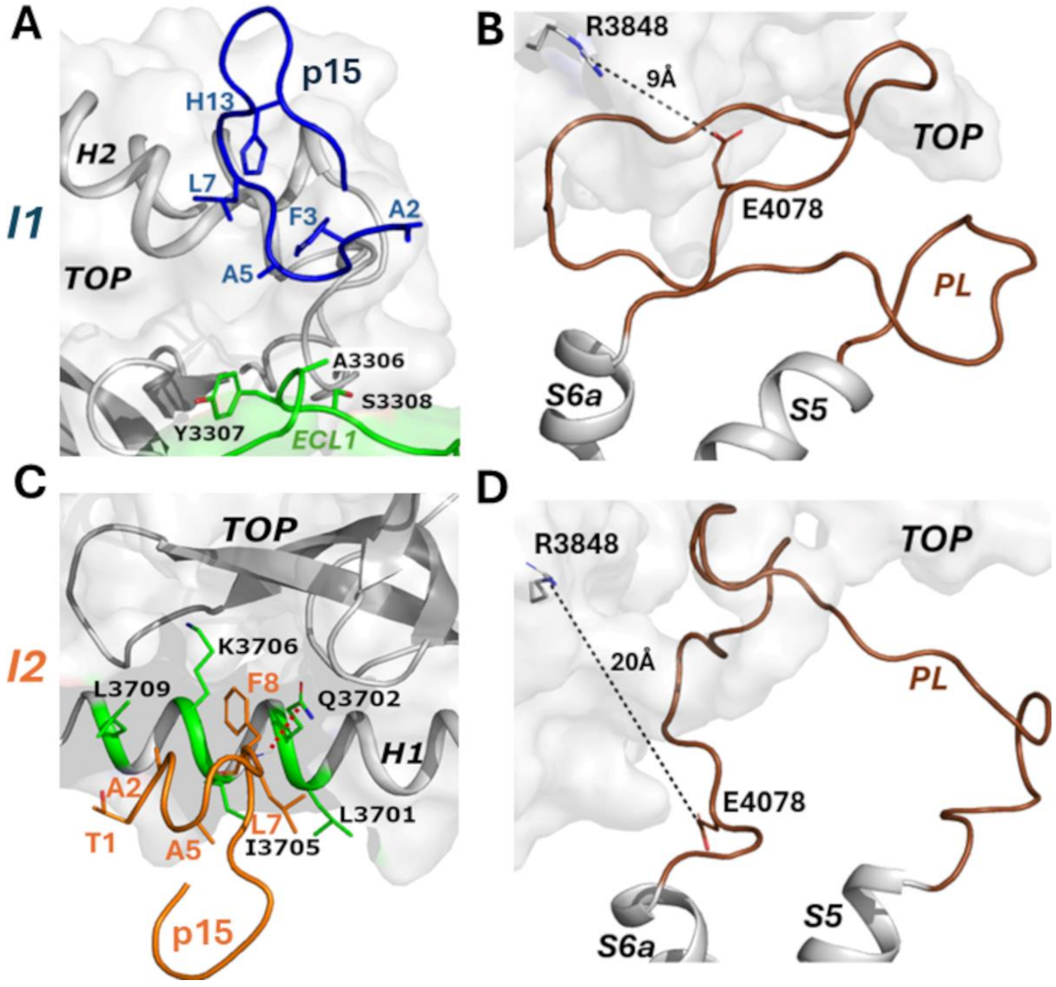

**Figure S10. Representative low-energy conformational states of p9, p15 and p17 binding to CTF-R3063C: (A)** Interactions observed between peptide-protein residues and **(B)** the distance between the TOP domain residue R3848 and pore loop (PL) residue E4078 observed in Intermediate 2 (I2) state of p9-bound CTF-R3063C. **(C)** Interactions observed between peptide-protein residues and **(D)** the distance between the TOP domain residue R3848 and pore loop (PL) residue E4078 observed in Intermediate 1 (I1) state of p15-bound CTF-R3063C. **(E)** Interactions observed between peptide-protein residues and **(F)** the distance between the TOP domain residue R3848 and pore loop (PL) residue E4078 observed in Intermediate 2 (I2) state of p15-bound CTF-R3063C. **(G)** Interactions observed between peptide-protein residues and **(H)** the distance between the TOP domain residue R3848 and pore loop (PL) residue E4078 observed in Intermediate 2 (I2) state of p17-bound CTF-R3063C.

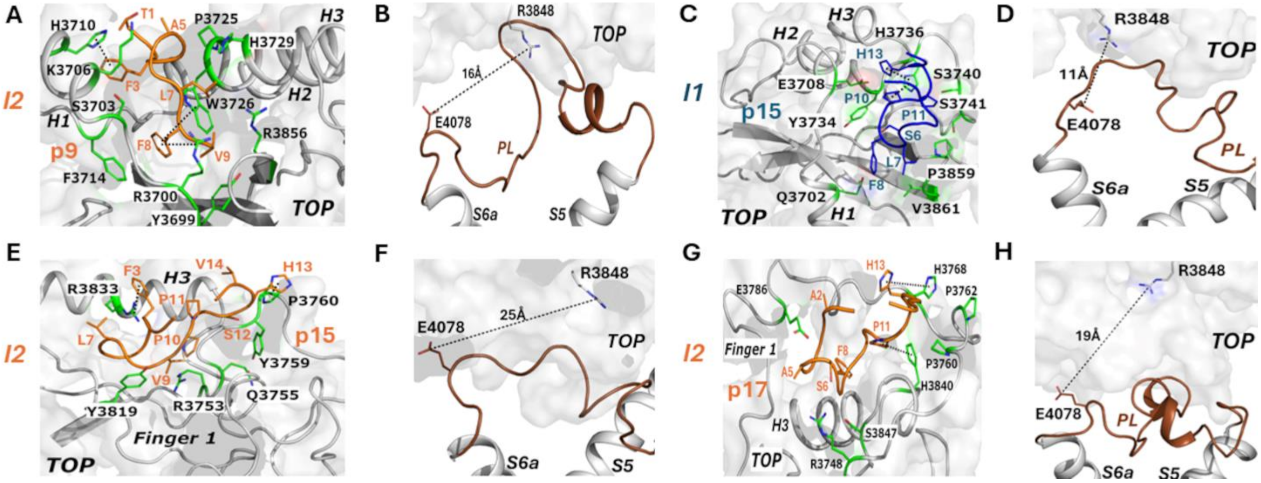

**Figure S11. Time courses of the number of contacts between the N-terminal region of Stalk (first 9 residues) and the TOP domain calculated from Pep-GaMD simulations of the: (A)** p9-bound CTF-G3052R, **(B)** p15-bound CTF-G3052R, **(C)** p17-bound CTF-G3052R, **(D)** p9-bound CTF-R3063C, **(E)** p15-bound CTF-R3063C, and **(F)** p17-bound CTF-R3063C systems, respectively. Residue contacts were calculated for heavy atoms with a distance cutoff of 4 Å.

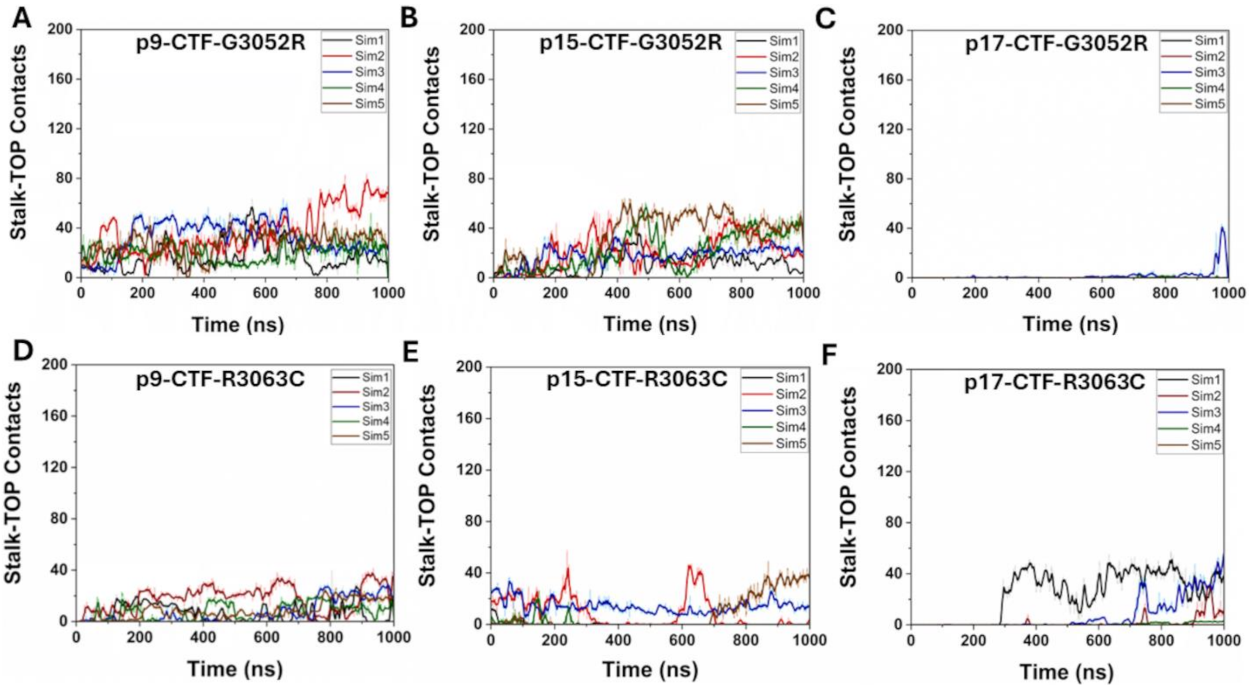

**Figure S12. Distortion of the TOP domain binding site in the presence of G3052R mutation in the p9-bound CTF-G3052R observed in Pep-GaMD simulations: (A)** Stalk formed one ionic interaction with the *H3* helix in the Intermediate 1 (I1) state of p9-bound CTF-G3052R. **(B)** Stalk formed polar interactions with the *H1* and *H2* helices in the Intermediate 2 (I2) state of p9-bound CTF-G3052R.

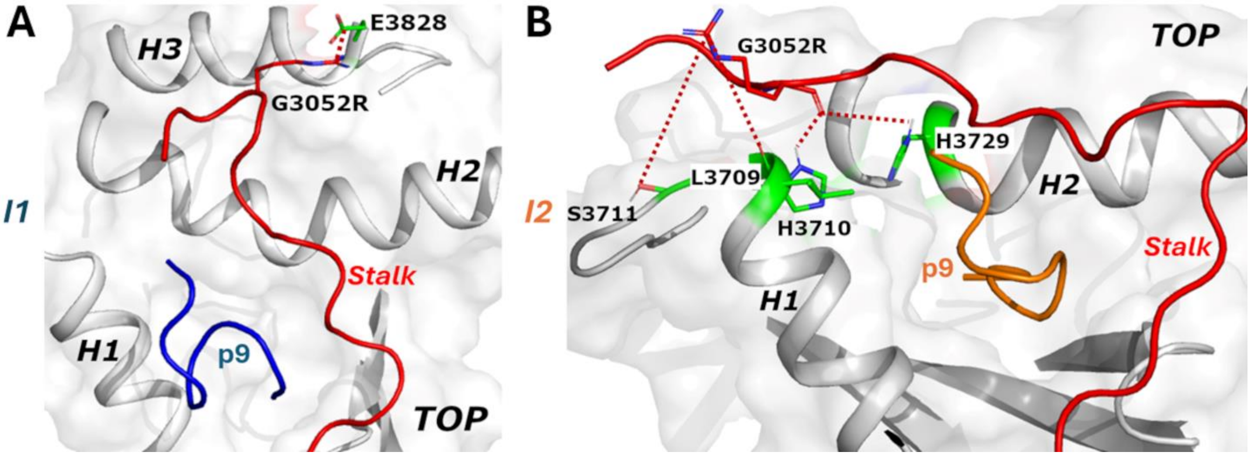

**Figure S13. Distortion of the TOP domain binding site in the presence of G3052R mutation in the p15-bound CTF-G3052R observed in Pep-GaMD simulations: (A)** Stalk formed one ionic interaction with the ECL1 loop in the Intermediate 1 (I1) state of p15-bound CTF-G3052R. **(B)** Stalk formed polar interactions with the ECL1 loop in the Intermediate 2 (I2) state of p15-bound CTF-G3052R.

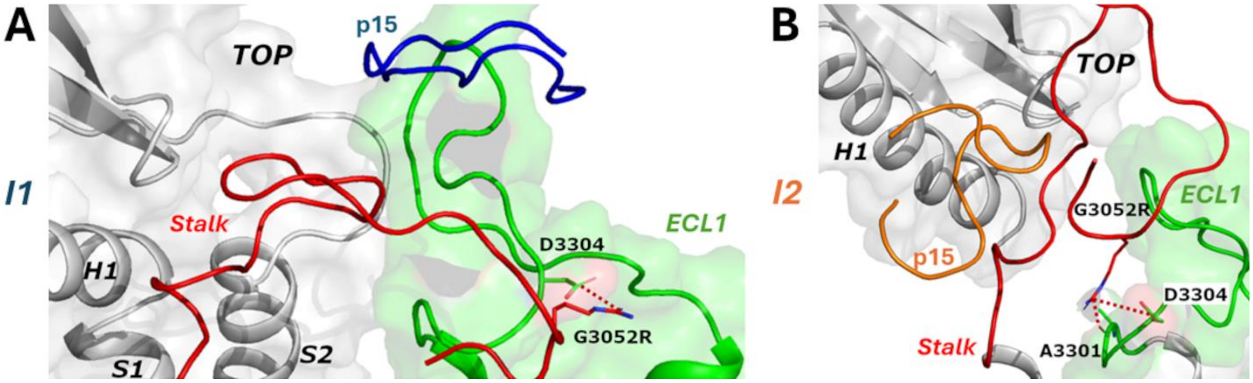

**Figure S14. TOP domain binding site maintained in Pep-GaMD simulations on p17-bound CTF-G3052R: (A-B)** p17 peptide stable binding on the TOP domain in the Intermediate 1 (I1) and Closed (C) states of p17-bound CTF-G3052R pushes the stalk outside with no interactions observed between the stalk and mutant CTF.

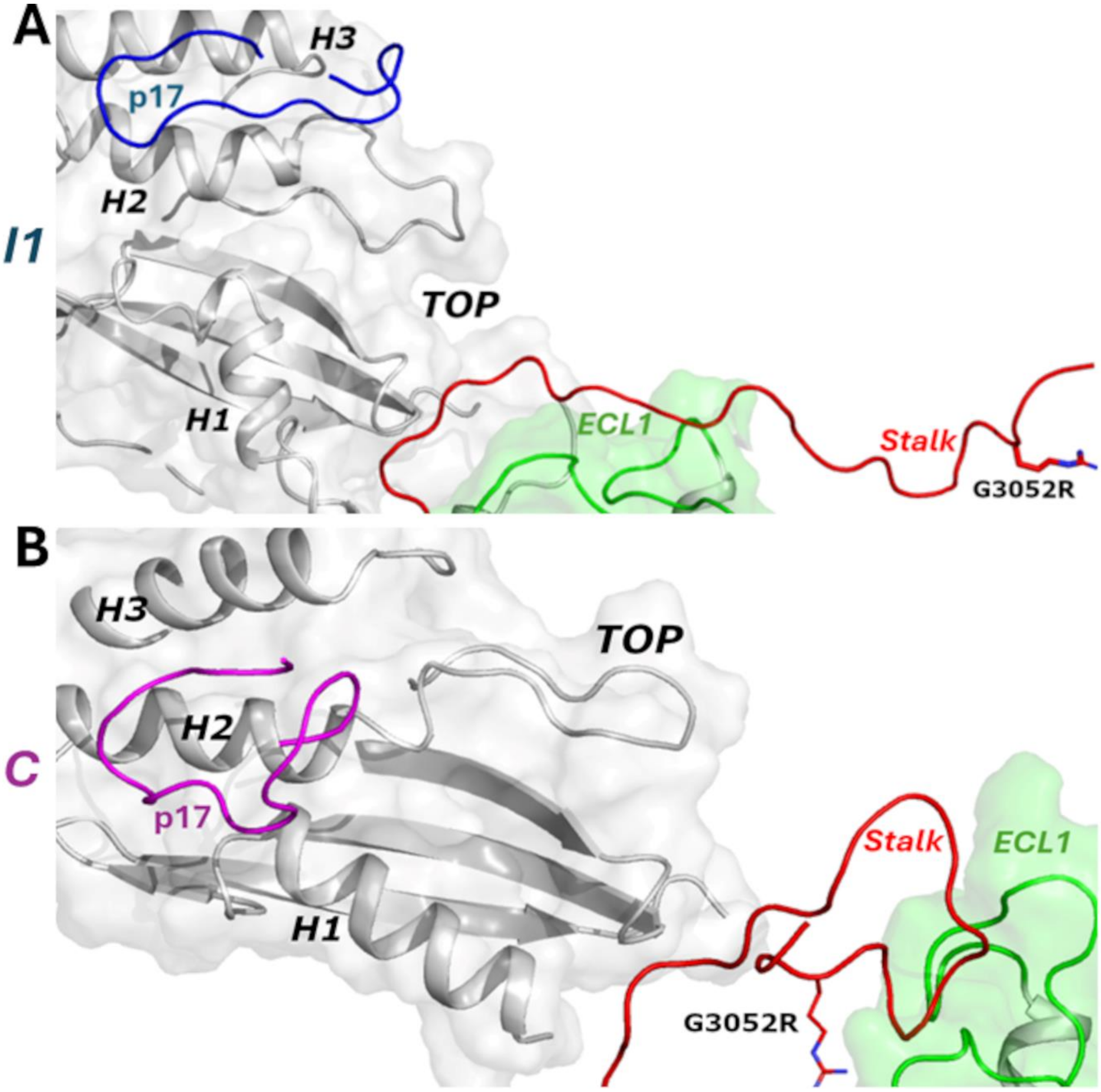

**Figure S15. TOP domain binding site maintained in Pep-GaMD simulations on p9-bound CTF-R3063C: (A)** Stalk binds to the *H3* helix in the TOP domain by formation of polar interactions in the Intermediate 2 (I2) state of p9-bound CTF-R3063C. **(B)** p9 peptide stable binding on the TOP domain in the Closed (C) state of p9-bound CTF-R3063C pushes the stalk outside with one ionic interaction observed between the stalk and the ECL1 loop in mutant CTF.

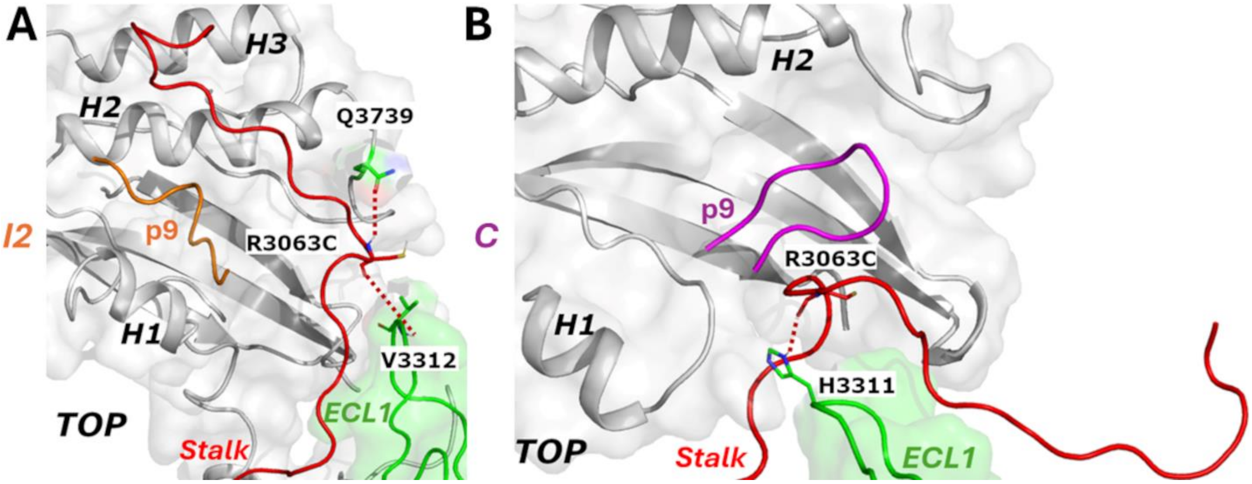

**Figure S16. TOP domain binding site maintained in Pep-GaMD simulations on p15-bound CTF-R3063C: (A)** p15 peptide stable binding on the TOP domain in the Intermediate 1 (I1) state of p15-bound CTF-R3063C pushes the stalk outside with no interactions observed between the stalk and mutant CTF. **(B)** Stalk forms one ionic interaction with the ECL1 loop in the Intermediate 2 (I2) state of p15-bound CTF-R3063C. **(C)** p15 peptide stable binding on the TOP domain in the Closed (C) state of p15-bound CTF-R3063C pushes the stalk outside with no interactions observed between the stalk and mutant CTF.

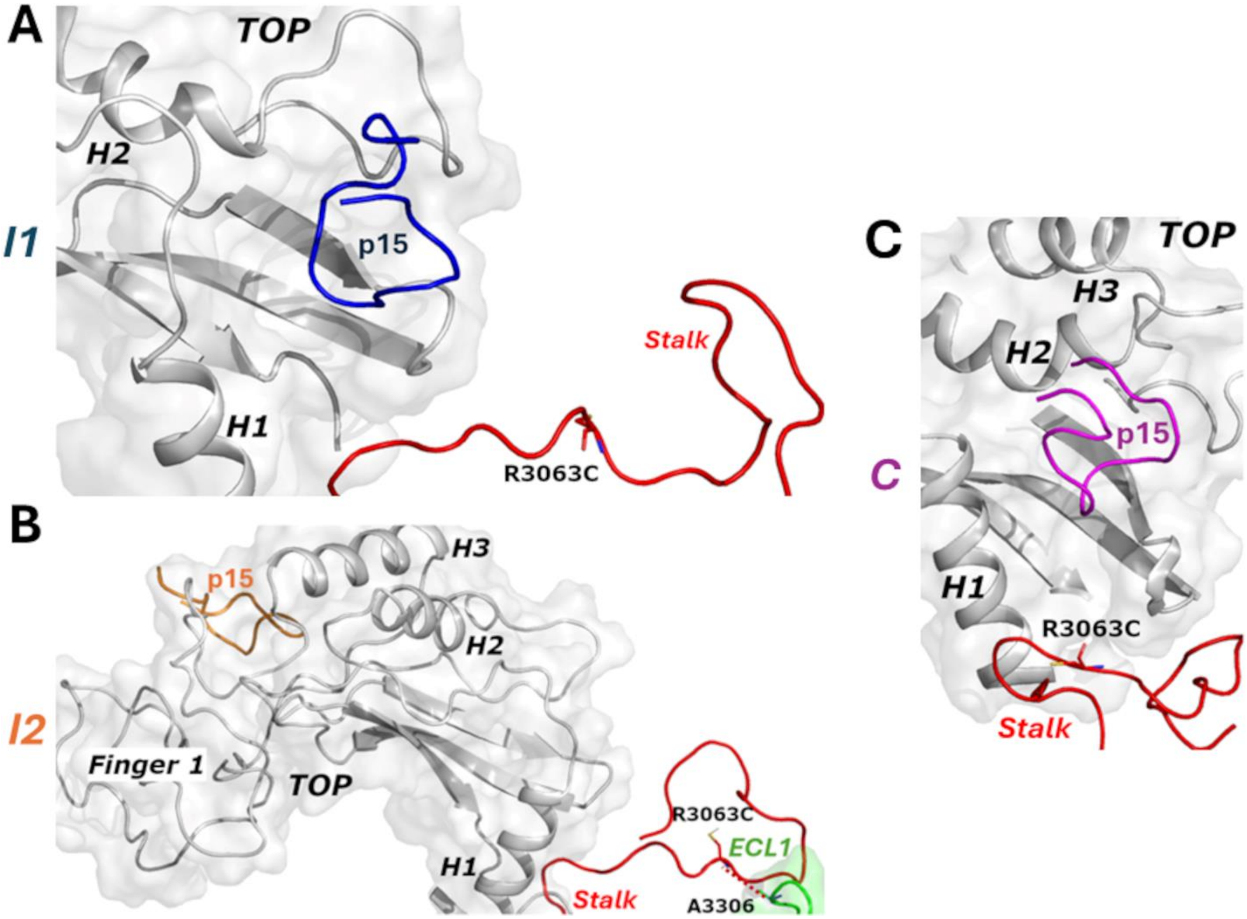

**Figure S17. TOP domain binding site maintained in Pep-GaMD simulations on p17-bound CTF-R3063C: (A)** Stalk forms polar interactions with the ECL1 loop in the Intermediate 2 (I2) state of p17-bound CTF-R3063C. **(B)** p17 peptide stable binding on the TOP domain in the Closed state (C) of p17-bound CTF-R3063C pushes the stalk outside with no interactions observed between the stalk and mutant CTF.

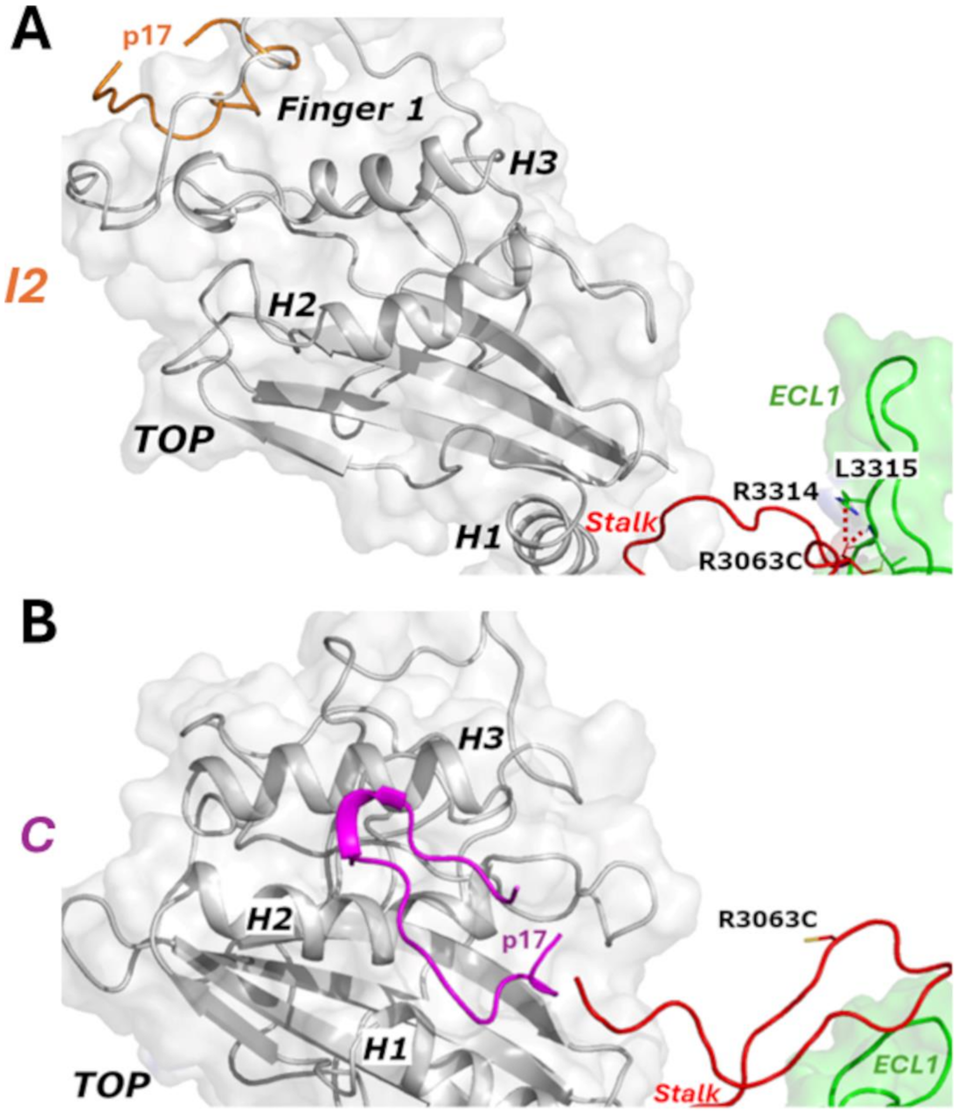

**Figure S18. Two-dimensional free energy profiles regarding the N3074-S3585 distances versus the Stalk-TOP contacts of the: (A)** p9-bound CTF-G3052R, **(B)** p15-bound CTF-G3052R, **(C)** p17-bound CTF-G3052R, **(D)** p9-bound CTF-R3063C, **(E)** p15-bound CTF-R3063C, and **(F)** p17-bound CTF-R3063C systems. The distances between the CG atom in N3074 and the OG atom in S3585 and the number of contacts between the N-terminal region of Stalk and the TOP domain were calculated from the Pep-GaMD simulations. Residue contacts were calculated for heavy atoms with a distance cutoff of 4 Å. Important low-energy conformational states are identified, including the "Closed," and "Open.

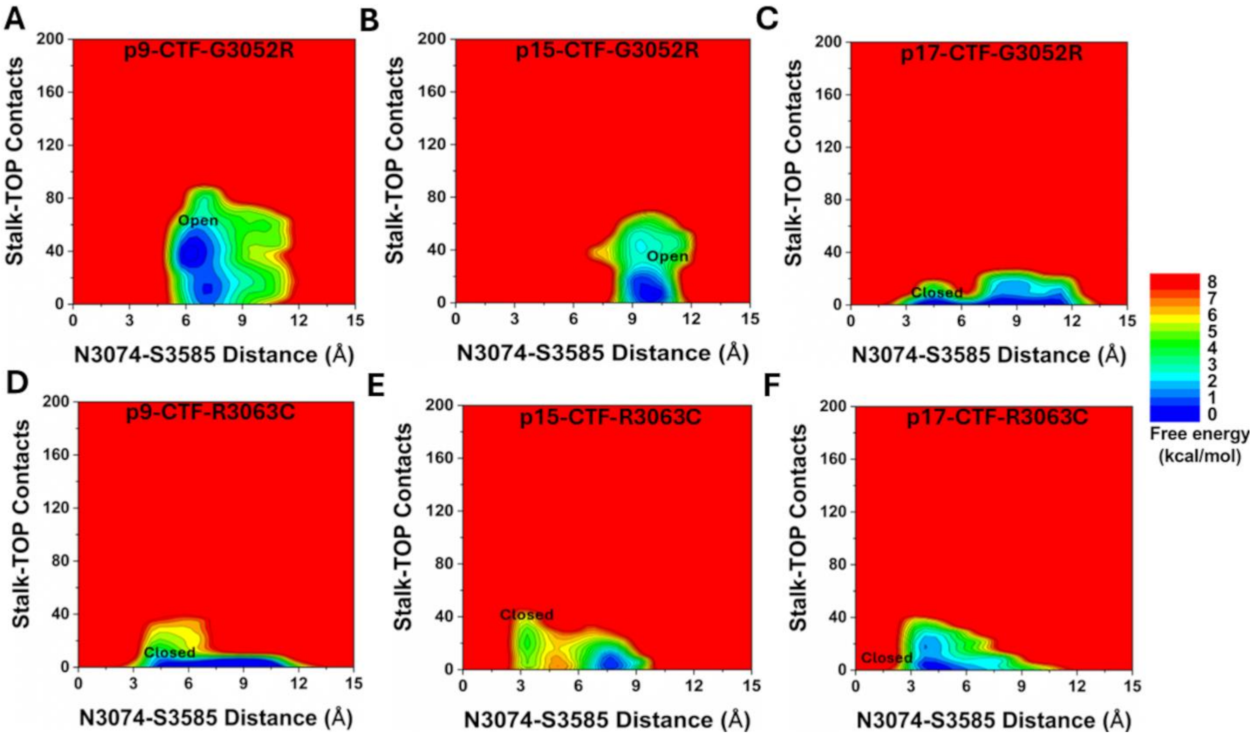

**Figure S19. Time course plots for the angle between the *H1* and *H2* helices for CTF mutants with p9, p15, and p17: (A)** Structural figure demonstrating an increase in the angle between the *H1* and *H2* helices for the active PC1 CTF[14] compared to the cryo-EM structure (PDB: 6A70)[15]. **(B-G)** Time courses of the angle between the *H1* and *H2* helices calculated from Pep-GaMD simulations of the **(B)** p9-bound CTF-G3052R, **(C)** p15-bound CTF-G3052R, **(D)** p17-bound CTF-G3052R, **(E)** p9-bound CTF-R3063C, **(F)** p15-bound CTF-R3063C, and **(G)** p17-bound CTF-R3063C systems, respectively.

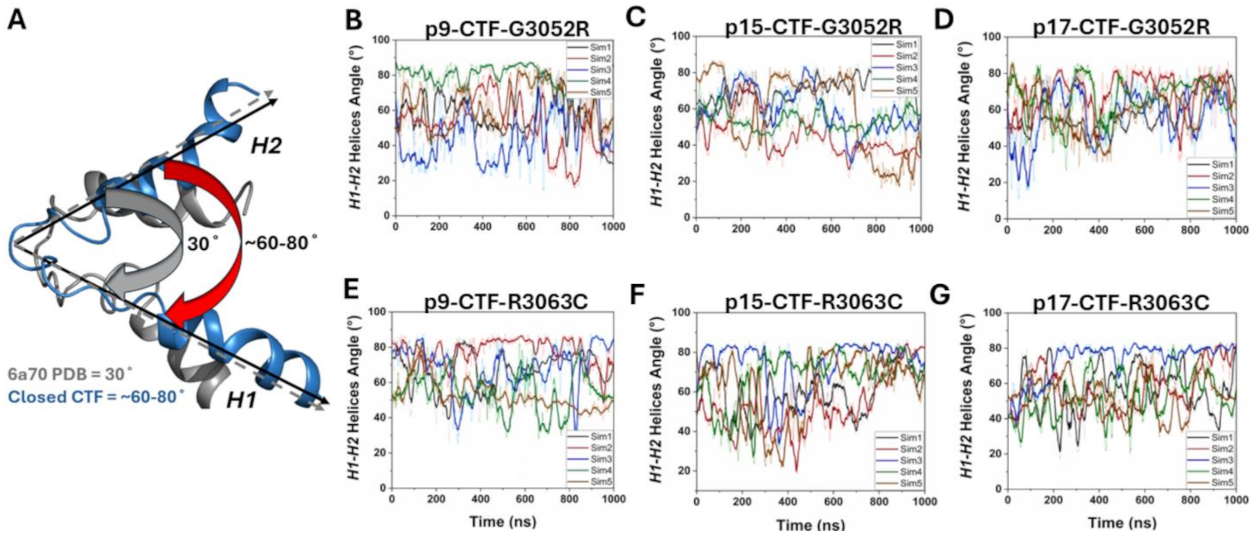

**Figure S20. Free energy profiles based on the angle between the *H1* and *H2* helices for CTF mutants with p9, p15, and p17: (A-F)** Two dimensional free energy profiles regarding the angle between the *H1* and *H2* helices and the R3848-E4078 distances calculated from Pep-GaMD simulations of the **(A)** p9-bound CTF-G3052R, **(B)** p15-bound CTF-G3052R, **(C)** p17-bound CTF-G3052R, **(D)** p9-bound CTF-R3063C, **(E)** p15-bound CTF-R3063C, and **(F)** p17-bound CTF-R3063C systems, respectively. Important low-energy conformational states are identified, including 'Open', 'Intermediate 2' (I2), 'Intermediate 1' (I1) and 'Closed'.

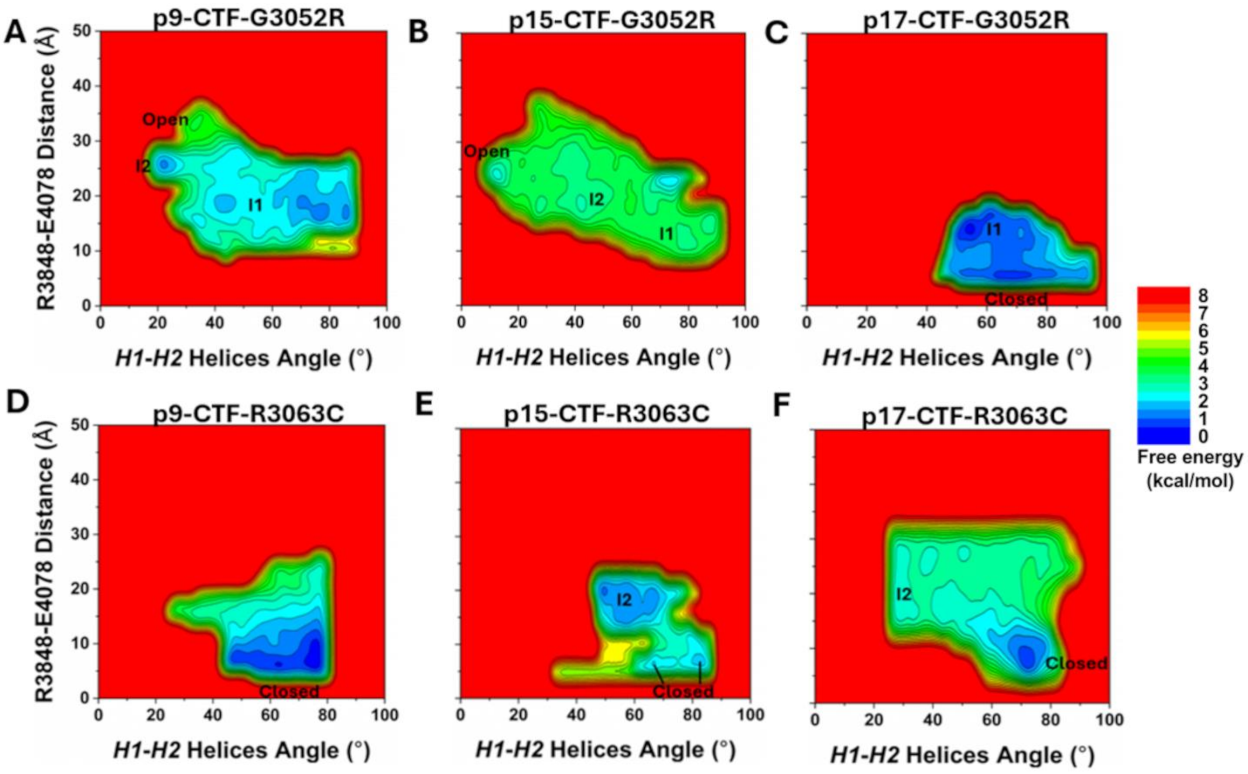

**Movie S1**: Representative movie for p9-bound CTF-G3052R system. PC1 protein is shown as light grey cartoon while stalk is shown as red cartoon and p9 peptide is shown as purple cartoon, respectively. For p9-bound CTF-G3052R mutant, the tethered stalk contacted the TOP domain more frequently, leading to an increased number of stalk-TOP residue contacts and formed polar interactions between R3052 and residues in the TOP domain.

**Movie S2**: Representative movie for p17-bound CTF-G3052R system. PC1 protein is shown as light grey cartoon while stalk is shown as red cartoon and p17 peptide is shown as purple cartoon, respectively. Binding of p17 stalk-derived peptide to the TOP domain prevented the stalk interaction with the TOP, leading to a decreased number of stalk-TOP residue contacts ($\leq 40$) and thereby inhibited any interactions between the residue R3052 and residues in the TOP domain.

**Movie S3**: Representative movie for p9-bound CTF-R3063C system. PC1 protein is shown as light grey cartoon while stalk is shown as red cartoon and p9 peptide is shown as purple cartoon, respectively. Tethered stalk showed higher fluctuations and mostly moved away from the TOP domain, leading to a decreased number of stalk-TOP residue contacts ($\leq 40$) and thereby inhibited any interactions between the residue C3063 and residues in the TOP domain.

#### 2. References

(1) Miao, Y.; Feher, V. A.; McCammon, J. A. Gaussian accelerated molecular dynamics: unconstrained enhanced sampling and free energy calculation. *Journal of chemical theory and computation* **2015**, *11* (8), 3584-3595.
(2) Miao, Y.; Sinko, W.; Pierce, L.; Bucher, D.; Walker, R. C.; McCammon, J. A. Improved reweighting of accelerated molecular dynamics simulations for free energy calculation. *Journal of chemical theory and computation* **2014**, *10* (7), 2677-2689.
(3) Hamelberg, D.; de Oliveira, C. A. F.; McCammon, J. A. Sampling of slow diffusive conformational transitions with accelerated molecular dynamics. *The Journal of chemical physics* **2007**, *127* (15).
(4) Burger, W. A.; Mobbs, J. I.; Rana, B.; Wang, J.; Joshi, K.; Gentry, P. R.; Yeasmin, M.; Venugopal, H.; Bender, A. M.; Lindsley, C. W. Cryo-EM reveals an extrahelical allosteric binding site at the M5 mAChR. *Nature communications* **2025**, *16* (1), 7046.
(5) Copeland, M. M.; Do, H. N.; Votapka, L.; Joshi, K.; Wang, J.; Amaro, R. E.; Miao, Y. Gaussian accelerated molecular dynamics in OpenMM. *The Journal of Physical Chemistry B* **2022**, *126* (31), 5810-5820.
(6) Joshi, K.; Wang, J.; Miao, Y. Enhanced Sampling of Biomolecular Interactions with Gaussian Accelerated Molecular Dynamics. *Computational Modeling of Biomolecular Interactions: Methods and Applications* **2026**, 493-516.
(7) Do, H. N.; Wang, J.; Joshi, K.; Koirala, K.; Miao, Y. Gaussian Accelerated Molecular Dynamics in Drug Discovery. *Computational Drug Discovery: Methods and Applications* **2024**, *1*, 21-43.
(8) Koirala, K.; Joshi, K.; Adediwura, V.; Wang, J.; Do, H.; Miao, Y. Accelerating molecular dynamics simulations for drug discovery. In *Computational drug discovery and design*, Springer, 2023; pp 187-202.
(9) Wang, J.; Miao, Y. Peptide Gaussian accelerated molecular dynamics (Pep-GaMD): Enhanced sampling and free energy and kinetics calculations of peptide binding. *The Journal of chemical physics* **2020**, *153* (15).
(10) Vanommeslaeghe, K.; MacKerell Jr, A. CHARMM additive and polarizable force fields for biophysics and computer-aided drug design. *Biochimica et Biophysica Acta (BBA)-General Subjects* **2015**, *1850* (5), 861-871.
(11) Duan, Y.; Wu, C.; Chowdhury, S.; Lee, M. C.; Xiong, G.; Zhang, W.; Yang, R.; Cieplak, P.; Luo, R.; Lee, T. A point-charge force field for molecular mechanics simulations of proteins based on condensed-phase quantum mechanical calculations. *Journal of computational chemistry* **2003**, *24* (16), 1999-2012.
(12) Eastwood, M. P.; Hardin, C.; Luthey-Schulten, Z.; Wolynes, P. G. Statistical mechanical refinement of protein structure prediction schemes: Cumulant expansion approach. *The Journal of chemical physics* **2002**, *117* (9), 4602-4615.
(13) Wang, Y.; Wang, Z.; Pavel, M. A.; Ng, C.; Kashyap, P.; Li, B.; Morais, T. D.; Ulloa, G. A.; Yu, Y. The diverse effects of pathogenic point mutations on ion channel activity of a gain-of-function polycystin-2. *Journal of Biological Chemistry* **2023**, *299* (5).
(14) Pawnikar, S.; Magenheimer, B. S.; Munoz, E. N.; Maser, R. L.; Miao, Y. Mechanism of tethered agonist-mediated signaling by polycystin-1. *Proceedings of the National Academy of Sciences* **2022**, *119* (19), e2113786119.

(15) Su, Q.; Hu, F.; Ge, X.; Lei, J.; Yu, S.; Wang, T.; Zhou, Q.; Mei, C.; Shi, Y. Structure of the human PKD1-PKD2 complex. *Science* **2018**, *361* (6406), eaat9819.
